# Basic interactions responsible for thymus function explain convoluted medulla shape

**DOI:** 10.1101/2024.09.29.615586

**Authors:** David Muramatsu, Henrik Weyer, Florian Gartner, Erwin Frey

## Abstract

The thymus is one of the most important organs of the immune system. It is responsible for both the production of T cells and the prevention of their autoimmunity. It comprises two types of tissue: the cortex, where nascent T cells (thymocytes) are generated; and the medulla, embedded within the cortex, where autoreactive thymocytes are eliminated through negative selection. In mice, the medulla exhibits a complex, convoluted morphology, which has raised the question of whether its form impacts its function. Intriguingly, experiments also reveal a reverse dependency: the interactions between medullary stroma and thymocytes shape the medullary structure. However, understanding the underlying mechanisms of medulla morphogenesis emerging from these interactions remains elusive. Here, we present a conceptual theoretical model which shows that central, experimentally verified signaling pathways suffice to shape the convoluted medullary structure. The mathematical analysis of the model explains the observed effects of chemotaxis on thymocyte localization, as well as the reported morphological changes resulting from the modulation of thymocyte production. Our findings reveal that the established cross-talk between medulla growth and negative selection of thymocytes not only regulates medullary volume but also orchestrates the morphology of the thymus medulla. This mechanism of structure formation robustly organizes the medulla in a way that accelerates thymocyte negative selection by improving their chemotactic migration into the medulla. Thereby, we identify a feedback between the function of the thymus medulla and its form. Our theoretical study motivates further experimental analysis of the spatial distribution of thymic cell populations and predicts morphological changes under genetic perturbations.

## I. INTRODUCTION

T cells are able to very specifically recognize proteins of pathogens and thus are central to the immune system. At the same time, self-reactive T cells that erroneously react to body protein can incite an autoimmune response. Consequently, the thymus, an organ in which new T cells are produced and self-reactive T cells are deleted, is critical for the development of an effective immune system. On an organ-wide scale, the thymus is made up of two functionally equivalent lobes. Each lobe consists of two microenvironments: an outer cortex and an inner, functionally different medulla; both microenvironments are built out of two types of epithelial cells, which are called cortical and medullary thymic epithelial cells (cTEC/mTEC) [1], Fig. 1*A*. In the cortex, nascent T cells, called thymocytes, develop from progenitor cells and have their ability to recognize protein checked (positive selection) [1]. Subsequently, thymocytes are chemotactically guided into the medulla where they are selected against reactivity to proteins they may find in the body’s tissues (negative selection).

**Figure 1.**
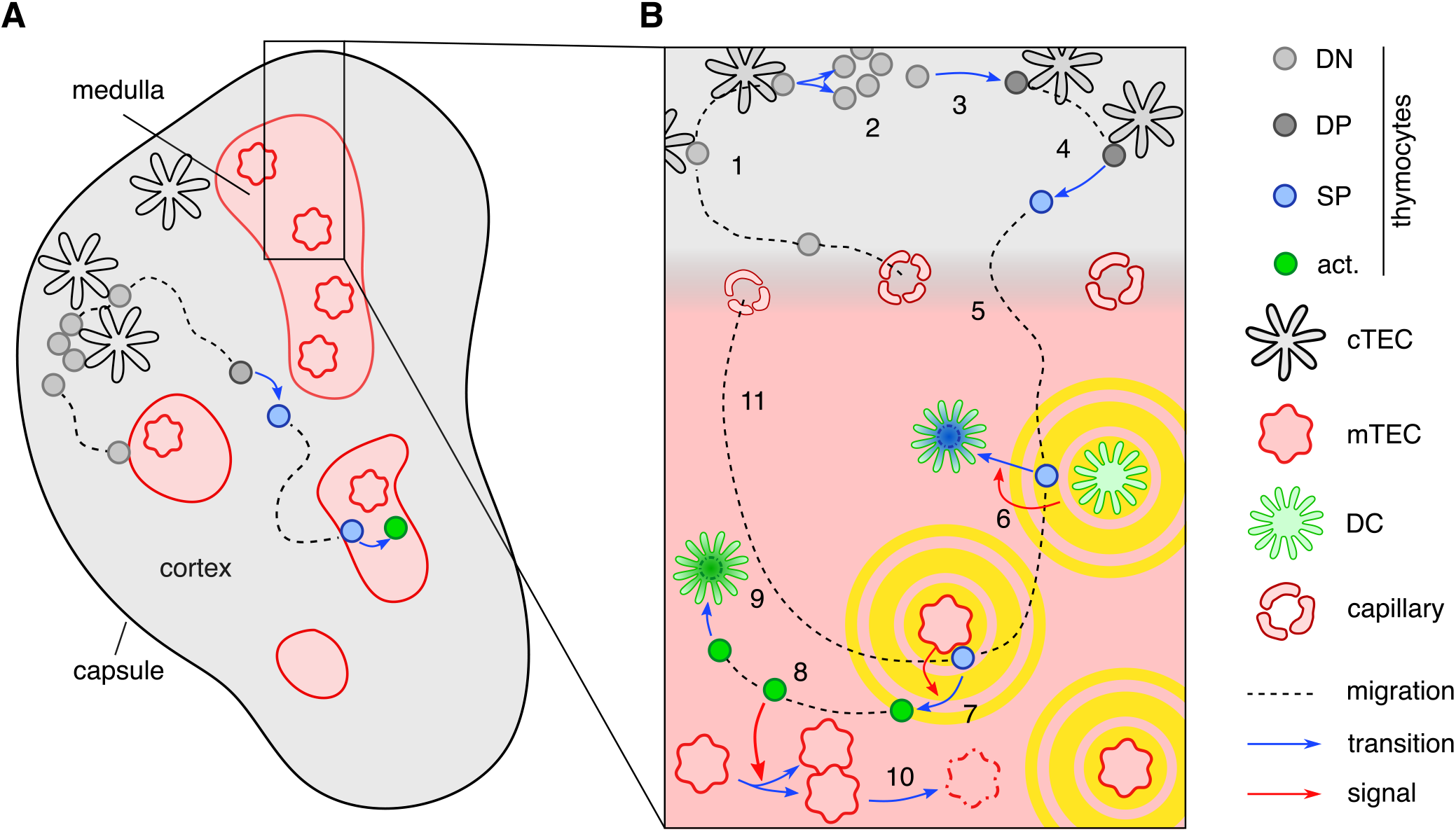
Thymocyte differentiation and thymic cross-talk. (*A*) The thymic tissue is separated into cortical (grey) and medullary (red shaded) regions, shown here for a cross-section through one lobe of the organ. Cortical stroma is formed predominantly by cTECs, while medullary stroma consists mainly of mTECs. (*B*) Early DN thymocytes enter the thymus through blood vessels. Subsequently, they migrate to the thymic capsule, which encloses the thymus (1). In the sub-capsular region, the DN thymocytes vigorously replicate (2). After replication, DN thymocytes differentiate into the CD4^+^ CD8^+^ double positive (DP) stage (3). These cells migrate to the deep cortex to be checked for basic functionality in a process called positive selection. DP thymocytes that successfully undergo positive selection transition (4) to the CD4^+^ CD8^-^ or CD4^-^ CD8^+^ single positive (SP) stage, and build up chemokine receptors which enables them to perform chemotaxis (5) into the thymus medulla. There, they are checked for self-reactivity by dendritic cells (DCs) (6) and medullary thymic epithelial cells (mTECs) (7). Self-reactive SP thymocytes that are found to be self-reactive by DCs are deleted by phagocytosis, whereas those that encountered their cognate antigen on mTECs enter an activated state, in which they can transmit differentiation and proliferation signals to mTECs and mTEC-precursors (8) before they are deleted by phagocytes (9). At the same time, both mTECs and DCs produce an array of chemoattractants that guide SP thymocytes to the medulla (circular yellow shading). The turnover of mTECs is high in the thymus medulla, such that mTEC death (10) is an important process in the thymus. Finally, thymocytes that have not encountered their cognate antigen in the medulla will exit the thymus after 4-5 days of migration inside medullary tissue (11).

Intriguingly, the medulla not only identifies self-reactive thymocytes but relies itself on self-reactive thymocytes for differentiation and growth signals [2–4]. This two-way communication between thymocytes and thymic epithelium has been termed thymic cross-talk [5, 6].

Thereby, the medulla obtains its shape (morphology) alongside the thymocyte selection process and does not pose a static, predefined environment. In mice, the resulting structure of the thymus medulla possesses an intricate convoluted morphology intertwined with cortical regions [7, 8]. This shape has been speculated to impact the efficiency of negative selection [7].

While the production [9] and selection [10] of thymocytes as well as the dynamics of the overall cTEC and mTEC populations [11] have been modeled numerically, the morphogenesis of the murine thymus medulla has not been investigated theoretically. Experimentally, the morphology of the medulla has been analyzed in two-dimensional slices and in its full three-dimensional structure [7, 8]. Interactions between thymocytes and epithelium have been analyzed on a mechanistic level [12, 13] as well as concerning their effects on thymus morphology [14, 15].

Here, we show how the thymus tissue can dynamically self-organize into a convoluted cortico–medullary structure based on only those interactions that are central to negative selection of thymocytes and medullary cross-talk. Our spatially explicit mathematical model predicts that these interactions lead to the segregation of the thymus into cortex and medulla through both chemotaxis-dependent and chemotaxis-independent mechanisms, which is consistent with experiments investigating the effects of suppressed thymocyte chemotaxis [16–22]. Furthermore, within the framework of our model, the morphology of the cortico-medullary pattern is set by both the volume fraction and an intrinsic length scale resulting from the interplay between thymocyte production and degradation. We make predictions to systematically test the connection between thymocyte dynamics and medullary structure and to recapitulate the phenotype of insular medullae in *H2-Aa*^*-/-*^ mice [7]. Mechanistically, we find that the medulla adapts its shape to the local supply of activated thymocytes. Thereby, the medulla self-organizes into an efficient cortex–medulla pattern, in which chemotaxis increases the effective degradation rate of self-reactive thymocytes compared to a thymus without cortex–medulla segregation.

## II. SPATIAL THYMIC CROSS-TALK MODEL

The thymus harbors a large number of interactions, many of which are realized redundantly. To provide the wider context of our model, we first describe the basic functionality of the thymus and those interactions important in thymocyte differentiation and cross-talk (Fig. 1B). For a more in-depth discussion of the interactions between thymocytes and thymic stroma, as well as the involved cells, we refer the interested reader to Refs. [12, 13, 23, 24]. We then introduce a reduced mathematical model, in which we focus on those interactions that are essential to medullary growth. Subsequently, we discuss our choice of the model parameters informed by experimentally known quantities.

### A. Thymocyte differentiation and medullary cross-talk

The central task of the thymus is to produce thymocytes and subject them to positive and negative selection. The thymocyte differentiation stages during this process are roughly characterized by the expression of two membrane proteins, CD4 and CD8. Early thymocytes in the CD4 and CD8 double negative (DN) stadium enter the thymus through blood vessels and migrate to the enclosing connective tissue (capsule) of the thymus (Fig. 1*B*). There, they proliferate and differentiate into double-positive (DP) thymocytes, which relocate deeper into the cortex and undergo positive selection [25]. During these two stages, each thymocyte develops a receptor to recognize specific proteins, called its cognate antigen, and checks the receptor for general functionality. After passing positive selection, thymocytes begin to build receptors for multiple chemoattractants—called chemokines—that are produced in the medulla, and the thymocytes transition to the CD4^+^CD8^-^ (CD4 SP) or to the CD4^-^CD8^+^ (CD8 SP) single-positive (SP) stage. The chemokines are produced by mTECs and dendritic cells (DC), and they form gradients pointing towards the medulla along which SP thymocytes orient their motion to localize to the medulla [20, 26–28].

In the medulla, thymocytes are presented with antigens derived from body protein by both mTECs and DCs. Self-reactive thymocytes thus can recognize their cognate antigen and subsequently enter an activated state. Activated CD4 and CD8 SP thymocytes undergo apoptosis and are cleared by phagocytosis within a few hours [29–31] or are converted into non-aggressive cell lines, such as regulatory T cells (only CD4 SP cells) [24]. CD4 SP thymocytes that are activated by antigen from an mTEC can stimulate mTECs and mTEC precursors into proliferation and differentiation [2–4, 14], and they are by far the largest source of the mTEC expansion signal [2, 3, 14]. Thymocytes that have not recognized a cognate antigen exit the thymus through blood vessels after 3-5 days [32] to build up the T-cell repertoire of the immune system.

### B. Reduced interaction network and mathematical model

To construct a mathematical model for the spatiotemporal dynamics of the cell populations, we adopt a coarse-grained perspective by organizing the pertinent cell populations and interactions according to their functions, as illustrated in Fig. 2A and discussed in further detail within the experimental context in the Supplementary Information (SI) Appendix.

**Figure 2.**
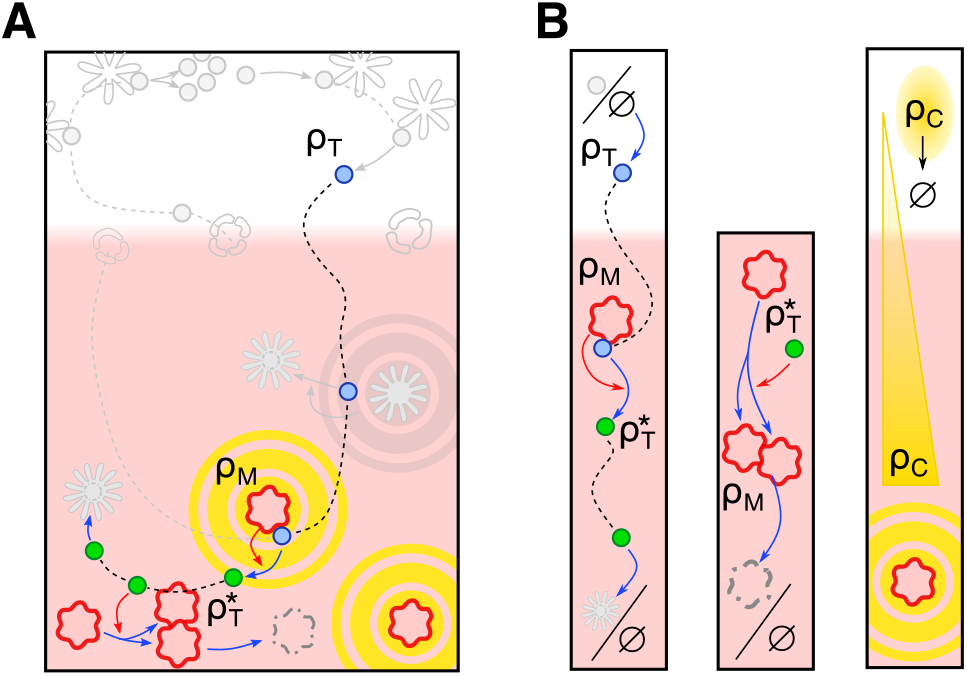
Reduced interaction network. (*A*) To describe the dynamics of medulla formation, we focus on interactions of (self-reactive) SP thymocytes with mTECs, with the same symbols as used in Fig. 1(*B*). These interactions include chemotaxis-driven migration of SP thymocytes *ρ*_T_ to the medulla, activation of self-reactive SP thymocytes, activated thymocyte-driven proliferation of mTECs *ρ*_M_, and the turnover of activated thymocytes 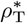 and mTECs. The processes not considered in the model are gray-shaded. (*B*) DP to SP differentiation is approximated as spatially homogeneous production of SP thymocytes in the cortex. Phagocytosis of activated self-reactive CD4 SP thymocytes is represented as a decay term (left column). Since we only consider self-reactive CD4 SP thymocytes, which do encounter their antigen on mTECs, egress of thymocytes and negative selection due to DCs are not included in the model. mTECs receive proliferation signals from activated thymocytes with an effective rate 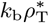 and die with a rate *k*_m_ (middle column). Chemokine emission with a rate *k*_e_ is limited to mTECs. Chemokines diffuse with diffusivity *D*_C_, and are degraded uniformly in the cortex and medulla with rate *k*_c_, thus forming gradients towards the medullary regions (right column). While the cells are depicted as particles, we model them in terms of local densities.

The medullary stroma comprises different subpopulations of mTECs [24], which we coarse-grain into a single effective mTEC population. Moreover, unless indicated otherwise we only model self-reactive CD4 SP thymocytes to give mTEC proliferation signals because these overwhelmingly stimulate mTEC proliferation [2, 3, 14]. Consequently, we will denote these self-reactive CD4 SP thymocytes simply as ‘(self-reactive) thymocytes’. Because DP thymocytes that differentiate into SP thymocytes are located in the cortex [25, 28, 33], we approximate the production of CD4 SP thymocytes by a homogeneous source in the cortex. Furthermore, since the exit rate of (self-reactive) thymocytes from the thymus is significantly lower than their activation rate, we disregard the export dynamics of the thymocytes. In summary, we include four local cell number densities and denote the densities of (self-reactive) thymocytes and activated thymocytes as *ρ*_T_ and 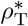, respectively, while referring to the densities of mTECs and chemokines as *ρ*_M_ and *ρ*_C_, respectively. The cortex is included implicitly as the regions of low mTEC density *ρ*_M_.

The thymocyte density *ρ*_T_ is increased through the production of SP thymocytes (from DP cells) with rate *p* in the cortex (Fig. 2*B* left). The spatial restriction to cortical regions is described using a smooth indicator function *ϕ*(*ρ*_M_) = (1 + (*ρ*_M_*/µ*)^2^)^−1^ with a density scale *µ*, which changes from *ϕ* ≈ 1 in cortical regions (*ρ*_M_ ≪ *µ*) to *ϕ* ≈ 0 in medullary regions with *ρ*_M_ ≫ *µ*. The conversion of *ρ*_T_ into activated thymocytes 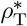 is due to activation by mTECs with a rate *k*_r_. Activated thymocytes undergo apoptosis and are eliminated by phagocytes [29– 31], which we model by a constant degradation term with rate *d*. Furthermore, we incorporate the chemotaxis of thymocytes along chemokine gradients using a Keller–Segel–type formulation [34–36], i.e., an interplay between random motion characterized by an effective diffusion coefficient *D*_T_ and directed motion with drift velocity v_C_ = *T χ*(*ρ*_C_) ∇*ρ*_C_ along gradients in the chemoat-tractant *ρ*_C_. Here, *T* quantifies the taxis strength and *χ*(*ρ*_C_) = *K*_OffOn_ · (*K*_OffOn_ + *ρ*_C_)^−2^ denotes a chemokine-dependent sensitivity resulting from receptor binding kinetics [37, 38]. Taken together, we have

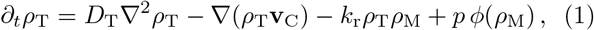

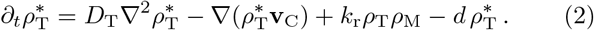

Before being degraded, activated thymocytes give proliferation signals to mTECs, which results in an effective proliferation rate 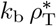 (second column of Fig. 2*B*). To account for the volume exclusion effects that limit mTEC growth beyond a certain density, we model mTEC proliferation in terms of logistic growth bounded by a maximum density *K* (carrying capacity). Together with the decay of mTECs at a rate *k*_m_, this results in the observed high mTEC-turnover rate [39]. As part of the stroma [25], mTECs are sedentary, and we only consider a weak random movement of the cells characterized by a small diffusion coefficient *D*_M_. The medullary tissue is thus described by

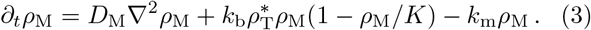

Finally, we describe the combined influence of all medullary chemokines on thymocytes using a single effective chemokine. This chemokine is produced with an effective rate *k*_e_ by medullary tissue *ρ*_M_ and diffuses in both cortex and medulla with an effective diffusion coefficient *D*_C_. Chemokines are degraded by interstitial proteases, and cellular uptake is facilitated by receptors such as CCRL1 [40]. For simplicity, we therefore include a uniform effective chemokine degradation rate *k*_c_, which gives

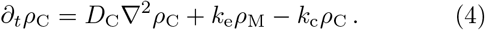

In summary, the equations for the dynamics of inactive and activated thymocytes, mTECs, and chemokines constitute a spatially explicit model for thymic cross-talk which can be viewed as a significantly extended Keller-Segel model [34]. The most important new features of the model include the two-state nature of the thymocytes and the presence of mTECs as secondary species. The proliferation of mTECs mediated by activated thymocytes and the mTEC’s production of a chemotactic field, which in turn controls the movement of the thymocytes organize the cells’ spatiotemporal dynamics. As outlined in the next section, the stimulated mTEC proliferation gives rise to a reaction–diffusion Turing-type instability [41] beyond the chemotaxis-induced instability of Keller– Segel models.

This spatial cross-talk model specifically focuses on the signaling between thymocytes and medullary tissue. It excludes mechanical effects due to cell growth that drive tissue flow. Therefore, we focus on the inner thymus region and simulate the inner thymic architecture within a fixed domain with periodic boundary conditions. The inclusion of the mechanics of tissue growth represents a fascinating challenge for future investigations.

The thymus medulla grows out of single cell ‘seeds’ with islands of monoclonal origin connecting to form larger medullary regions [42]. We therefore initialize the simulations with 300 small medullary regions [42] scattered randomly throughout the simulated volume.

### C. Choice of model parameters

We briefly detail the parameters used in the numerical analysis of the model, with a more detailed discussion of the relevant literature in the SI Appendix Sec I.B. Direct measurements give experimental values for the diffusivity *D*_T_ of thymocytes [43], the death rate *d* of activated thymocytes [30, 31], and the mTEC decay rate *k*_m_ [44]. The turnover of self-reactive thymocytes has been measured by [45], from which we estimate the production rate of self-reactive thymocytes.

The time scale of antigen recognition has been probed experimentally in the context of selection against densely [30] as well as sparsely presented antigen [31, 46]. We orient the thymocyte activation rate on these measurements, although additional measurements on the fraction of self-reactive T cells in the periphery suggest potentially longer time scales of antigen recognition [47, 48], as detailed in the SI Appendix Sec. I.

The mTEC carrying capacity *K* and the strength of mTEC proliferation, *k*_b_, are not known. We choose those parameters such that the medullary volume [7, 49], the total mTEC numbers [49], and the turnover of mTECs match the experimentally measured values.

Since we model a single effective chemokine, the rates for chemokine production and decay, *k*_e,c_, the binding affinity *K*_OffOn_ of chemokine to its receptor, the chemokine diffusivity *D*_C_, and the chemotaxis strength *T* are effective parameters that are not fixed by experimental values. Where possible, we orient the parameter choice along results on the chemokine signaling axis CCL21-CCR7, as it is the most important chemokine signal for medullary morphology [16, 50]. Unless stated otherwise, the simulated time corresponds to approximately 12 weeks. The standard parameter values chosen for the wild-type (wt) case are given in Tab. I.

**Table I.** Parameter values for the simulation reflecting wild-type conditions. Deviating parameter values are stated along-side the corresponding figures.

| parameter | symbol | value | unit |
| --- | --- | --- | --- |
| chemokine diffusivity | $D_C$ | $2.34 \cdot 10^4$ | $\mu\text{m}^2 \text{h}^{-1}$ |
| thymocyte random motility | $D_T$ | $2.4 \cdot 10^4$ | $\mu\text{m}^2 \text{h}^{-1}$ |
| taxis strength | $T$ | $10^5$ | $\mu\text{m}^2 \text{h}^{-1}$ |
| mTEC random motility | $D_M$ | 3 | $\mu\text{m}^2 \text{h}^{-1}$ |
| receptor binding constant | $K_{\text{OffOn}}$ | 5 | nM |
| thymocyte production | $p$ | $2.5 \cdot 10^{-6}$ | $\mu\text{m}^{-3} \text{h}^{-1}$ |
| mTEC density scale | $\mu$ | $10^{-4}$ | $\mu\text{m}^{-3}$ |
| thymocyte deletion | $d$ | 1 | $\text{h}^{-1}$ |
| thymocyte conversion | $k_r$ | $1.5 \cdot 10^3$ | $\mu\text{m}^3 \text{h}^{-1}$ |
| mTEC birth rate | $k_b$ | $3 \cdot 10^3$ | $\mu\text{m}^3 \text{h}^{-1}$ |
| mTEC carrying capacity | $K$ | $7.5 \cdot 10^{-4}$ | $\mu\text{m}^{-3}$ |
| mTEC decay | $k_m$ | $5 \cdot 10^{-3}$ | $\text{h}^{-1}$ |
| chemokine emission | $k_e$ | 0.014 | $\text{M} \mu\text{m}^3 \text{h}^{-1}$ |
| chemokine decay | $k_c$ | 10 | $\text{h}^{-1}$ |
| system size | $L$ | 3 | mm |
| simulated time | $t_{\text{max}}$ | $2 \cdot 10^3$ | h |

## III. RESULTS

On the basis of our spatial thymic cross-talk model (Eqs. 1–4), we now analyze the spatiotemporal dynamics of the densities of thymocytes, mTECs, and chemokines as a function of the kinetic parameters. Figure 3 shows the (quasi-)steady-state distribution of the mTEC density *ρ*_*M*_ and highlights different medulla morphologies that fall within the scope of our model. This includes sperical islets, ‘swiss-cheese’ morphologies, and convoluted structures. The medullary region (*ρ*_M_ *> µ*) for a parameter set within the physiological range (Tab. I) is shown in Fig. 3*A*. It displays a convoluted medullary morphology that matches the phenotype found in experiments on wild type (wt) mice [7, 8]. This qualitative phenomenology persists even for a ten-fold increase in thymocyte production and degradation (Fig. 3*B*). Choosing a lower thymocyte production rate results in insular medullae (Fig. 3*C*), and increasing it leads to a ‘swiss-cheese’-like medullary configuration (Fig. 3*D*). Both medullary morphologies match those previously found in ablation experiments [7, 51, 52].

**Figure 3.**
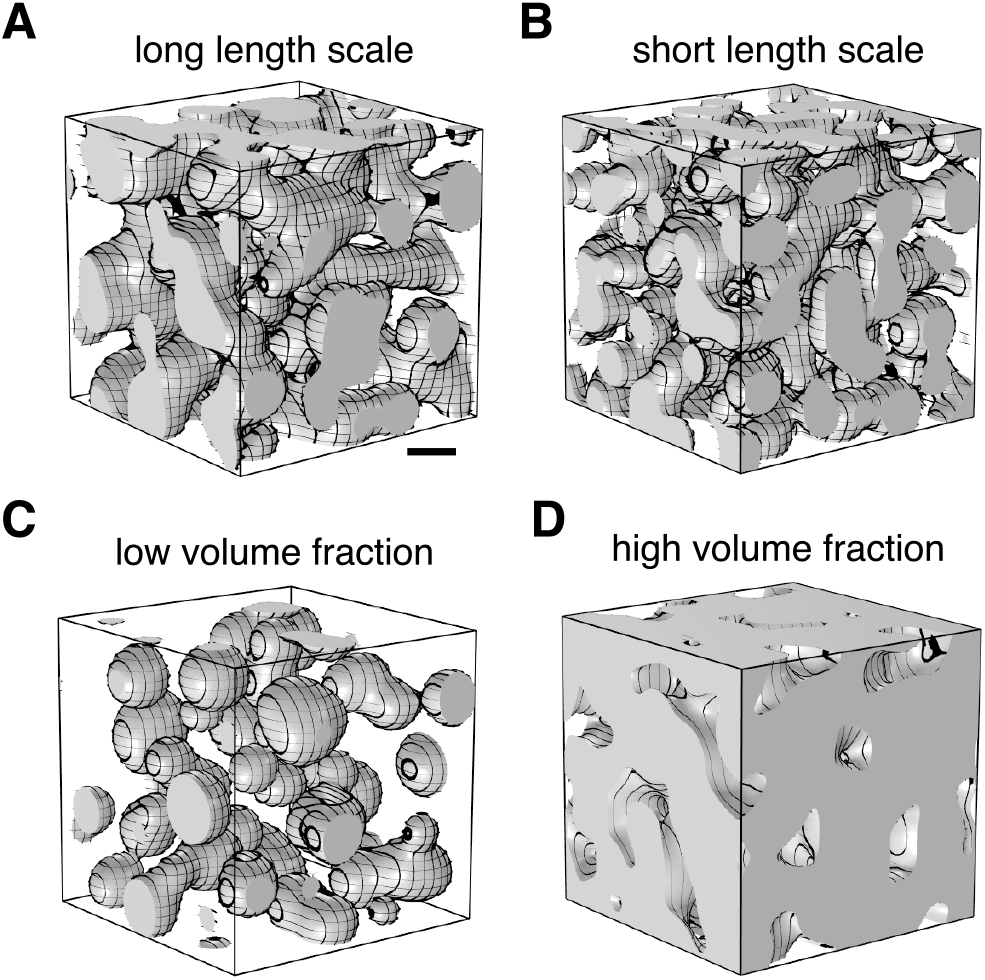
Medulla morphology: (Quasi-)steady-state densities *ρ*_M_ (*t* = 2000h). The spatial cross-talk model produces a *convoluted medullary* morphology for the parameter set Tab. I chosen within physiological ranges (*A*) and (*B*) for ten-fold increased thymocyte production (*p* = 2.5 · 10^−5^µm^−3^ h^−1^) and degradation (*d* = 10h^−1^). For a lower production rate *p* = 10^−6^µm^−3^ h^−1^, the model displays *insular medullae* (*C*) and for a heightened production rate *p* = 7 · 10^−6^µm^−3^ h^−1^, the medullae coalesce to a *‘swiss-cheese’* morphology. The morphology depends on the characteristic length scale set by the thymocyte production-degradation dynamics, as well as the medullary volume fraction *φ*_M_ = ⟨*θ*(*ρ*_M_ − 10^−4^µm^−3^)⟩, the fraction of space inhabited by the medulla. At low volume fraction *φ*_M_, the insular medullae (*C*: *φ*_M_ = 0.18) form, which elongate to form tubes, then connect to form the convoluted morphology (*A*: *φ*_M_ = 0.41, *B*: *φ*_M_ = 0.4), and finally coalesce to form the ‘swiss-cheese’ morphology (*D*: *φ*_M_ = 0.83) for increasing medullary volume fraction. Regions with *ρ*_*M*_ *> µ* are colored gray, scale bar: 500µm, volume side length *L* = 3mm. The simulation employs periodic boundary conditions and is initialized with 300 small, uniformly distributed medullary regions of width *w* = 60µm.

In the following sections, we dissect the mechanisms driving the emergence of the different morphologies by analyzing our model for variations in three key kinetic parameters: the chemotaxis strength *T*, the thymocytes production rate *p*, and the degradation rate *d* of activated thymocytes. Importantly, all of the above parameters can be varied through genetic perturbations; see e.g. Refs. [12, 15, 53].

### A. Pattern-forming instabilities

The spatiotemporal dynamics described by the spatial thymic cross-talk (STCT) model (Eqs. 1–4) exhibit a segregation into cortex and medulla. To gain initial insights into the dominant processes underlying the segregation, we investigate the stability of spatially uniform distributions of cell densities. In the model, two distinct types of instabilities contribute: (i) a chemotaxis-induced instability as in the Keller-Segel model, and (ii) a growth–induced (Turing) instability characterized by positive feedback in mTEC growth coupled to thymocyte diffusion. Both instabilities yield a band of unstable Fourier modes, but the mechanisms underlying the instabilities are qualitatively different.

The Keller-Segel-type instability is driven by chemotaxis accumulating thymocytes in regions with increased mTEC densities. As the (activated) thymocytes stimulate further mTEC proliferation, and thus increase the local chemoattractant production, this accumulation self-amplifies. The mechanism underlying the growth-induced instability is independent of chemotaxis. Instead, the mechanism is linked to the positive feedback in mTEC growth: mTECs activate thymocytes, and reciprocally, activated thymocytes stimulate mTEC proliferation. At fixed thymocyte density *ρ*_T_, this leads to a bistability in the mTEC dynamics between a state with low densities of both mTEC and activated thymocytes and a state with both species at high densities.

The lateral instabilities caused by both mechanisms separately, as well as their combination in the full model are summarized in a stability diagram changing the chemotaxis strength *T* and the mTEC proliferation rate *k*_b_ (Fig. 4*A*). The individual mechanisms are isolated by constructing two reduced models in which either the effective bistability in mTEC turnover (Fig. 4*A*, orange-shaded area) or chemotaxis is suppressed (fixing *T* = 0; Turing-unstable, blue-shaded region). Both instabilities synergize in the full model (gray-shaded; see SI Appendix Sec. II).

**Figure 4.**
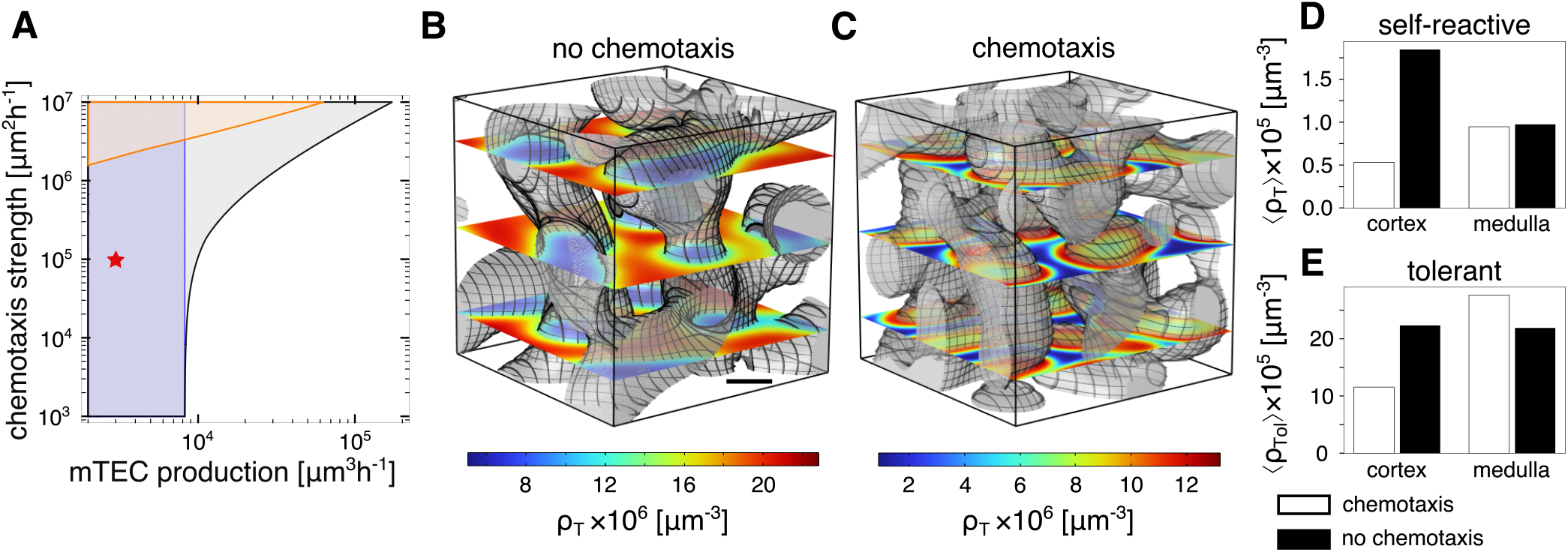
Cortex–medulla segregation and thymocyte localization. Cortex and medulla segregation is driven by chemotaxis and thymocyte selection, however efficient medullary localization of thymocytes relies on chemotaxis. (*A*) Linear stability analysis (LSA) shows that the model spontaneously segregates into cortex and medulla (gray-shaded domain) independently of chemotaxis at low mTEC proliferation rates *k*_b_ (blue-shaded box), while a chemotaxis-driven instability enlarges this regime towards larger proliferation rates *k*_b_ at high chemotaxis strength (orange-shaded triangle). The instability region for an approximative model that lacks the bistability in the mTEC dynamics is shown in the orange-shaded triangle. The red star denotes wt parameters. Without (*B, T* = 0) and with chemotaxis (*C*, wt parameters), the model shows a convoluted medullary morphology. However, in the wt scenario, self-reactive thymocytes localize efficiently to the medulla, while the self-reactive thymocytes show an inverted density ratio when chemotaxis is not present (*D*). Tolerant thymocytes accumulate in the medulla through chemotaxis but are almost evenly distributed when chemotaxis is absent (*E*). Scale bar: 500µm, rates are chosen as detailed in Tab. I.

The ability of thymic tissue to re-organize into a defined medulla and cortex after a complete disruption of the three-dimensional organization in reaggregate thymic organ cultures [54] (see for instance [55, 56]) indicates that the segregation of medulla and cortex does not rely on external cues but rather is a self-organized thymocyte-dependent process.

Furthermore, it has been found that the ablation of different chemotaxis signals, and even double-knockouts of chemokine receptors, do not abolish the segregation of cortex and medulla [16–22]. Thus, a chemotaxis-independent pattern formation mechanism is plausible.

In our model, the medullary morphology does not qualitatively change when chemotaxis vanishes. However, it has been shown that the CCL21 chemokine signal does influence medullary morphology with a knockout of either the chemokine or the chemokine receptor producing smaller, less connected medullae [16–18] with lower overall volume [50]. This effect may occur because ablating the CCR7-CCL21 signaling pathway not only influences the localization of thymocytes, but also lowers DN and DP thymocyte numbers[16, 17], possibly leading to a decreased production rate of SP thymocytes. As we explore in Sec. III C, a reduced thymocyte production can change the medullary morphology towards insular medullae.

### B. Chemotaxis impacts thymocyte localization

Discovering that chemotactic signaling is not a prerequisite for nontrivial medulla morphologies, our subsequent aim was to discern its influence on the spatial arrangement of mTECs and thymocytes. While the chemotaxis-induced instability is driven by the colocalization of thymocytes and mTECs, the growth-induced instability only includes the net degradation of thymocytes in medullary regions and net production in cortical regions. This suggests that in this case, self-reactive thymocyte densities are higher where mTECs are absent, and vice versa. We indeed observe this difference in simulations of the spatial thymic cross-talk model in the physiological parameter regime in the presence and absence of chemotactic signaling; compare Fig. 4*B* and *C*. Importantly, the experimentally observed ‘arrest’ of SP thymocytes in the cortex [50] is not due to a confinement of thymocytes to the cortex in our model. Rather, it is due to an activation and subsequent degradation process that is fast compared to the time scale of thymocyte random motion into the medulla.

Experimental evidence suggests that self-tolerant (pigeon cytochrome c-specific [57]) thymocytes show different localization patterns compared to self-reactive thymocytes when CCL21-driven chemotaxis is ablated [50]. To assess if our model can replicate this observation, we include the self-tolerant (CD4) thymocytes as an additional species 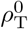. As these thymocytes do not participate in cross-talk [2, 3], they do not influence the dynamics and morphology of the other species. However, the morphology developing in the spatial thymic cross-talk model determines how these self-tolerant thymocytes distribute. Analogously to the self-reactive thymocytes, we model the self-tolerant thymocytes to be produced in the cortex with a rate 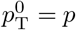 ([45], see SI Appendix Sec. I.B.2) and to possess the same motility as self-reactive thymocytes. After a dwell time of approximately five days [32], tolerant thymocytes exit the thymus in a process that is largely independent of their stay in the medulla [50]. This process is modeled by a homogeneous degradation rate *k*_ex_ = (5d)^−1^. The tolerant thymocyte population is thus described by

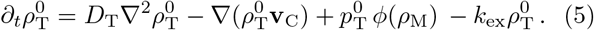

For completeness, we also add the thymus-exit term for the self-reactive thymocytes to the extended model for all simulations in this section.

With chemotaxis (parameters given in Tab. I), the self-reactive and self-tolerant thymocytes are concentrated in the medulla in the extended model Eqs. (1-5) with a medulla-to-cortex ratio of 2:1 for self-reactive thymocytes and of 3:1 for self-tolerant thymocytes (Fig. 4*C* - *E*). This agrees qualitatively with experimental results in wild-type mice yielding even higher ratios [26, 28, 50]. In the absence of chemotaxis (*T* = 0), self-reactive cells show a roughly 3:5 medulla-to-cortex density ratio in the simulation in qualitative accordance with the results for

CD4 SP in *Ccr7*^*-/-*^ mice [18, 50], in which the T cell-borne chemokine receptor CCR7 is ablated (Fig. 4*C,D*). At the same time, tolerant thymocytes show nearly a 1:1 density ratio between medulla and cortex in chemotaxis-deficient conditions (Fig. 4*E*), because the time they need to traverse the cortex–medulla pattern by diffusion is far lower than the time scale of their exit from the thymus (SI Appendix Sec. I.B.16). This uniform distribution of tolerant thymocytes is in accordance with the results in *Ccr7*^*-/-*^ and *plt* mice [50], in which the ligands of CCR7 are ablated.

Furthermore, we observe that the self-reactive thymocytes are mostly present at the interface of the medulla (cortico-medullary junction), and they fall off towards the interior of the medulla (Fig. 4(*C*), SI Appendix Fig. S2*A*), whereas self-tolerant thymocytes are distributed homogeneously throughout the medulla (SI Appendix Fig. S2*B*). Experiments that observe the full spatial distribution of self-reactive and self-tolerant thymocytes are essential for testing our prediction of medulla-interior structuring of the different thymocyte species. If our prediction holds, this could have interesting down-stream effects on the distribution of different types of mTECs when thymocyte signaling for mTEC differentiation into sub-species is taken into account, see e.g., Ref. [4].

### C. Thymocyte production and degradation govern the morphology and intrinsic length scale of the medulla

The amount of activated thymocytes dictates the rate at which mTECs are built, as they stimulate mTEC proliferation. Thus, in our spatial thymic cross-talk model, a lower amount of thymocytes—induced by a decrease in thymocyte production (rate *p*) or an increase in the degradation of activated thymocytes (rate *d*)—results in a reduced total mTEC number. As a consequence of this overall reduction in the population of activated thymocytes, the volume fraction of the medulla is reduced and one obtains an insular morphology. Opposite perturbations of the rates result in increased overall mTEC proliferation and a Swiss-cheese-like morphology. Reduced two-dimensional numerical simulations (parameters given in Tab. I) show this behavior for a large range of the rates *p* and *d* (Fig. 5*A*).

**Figure 5.**
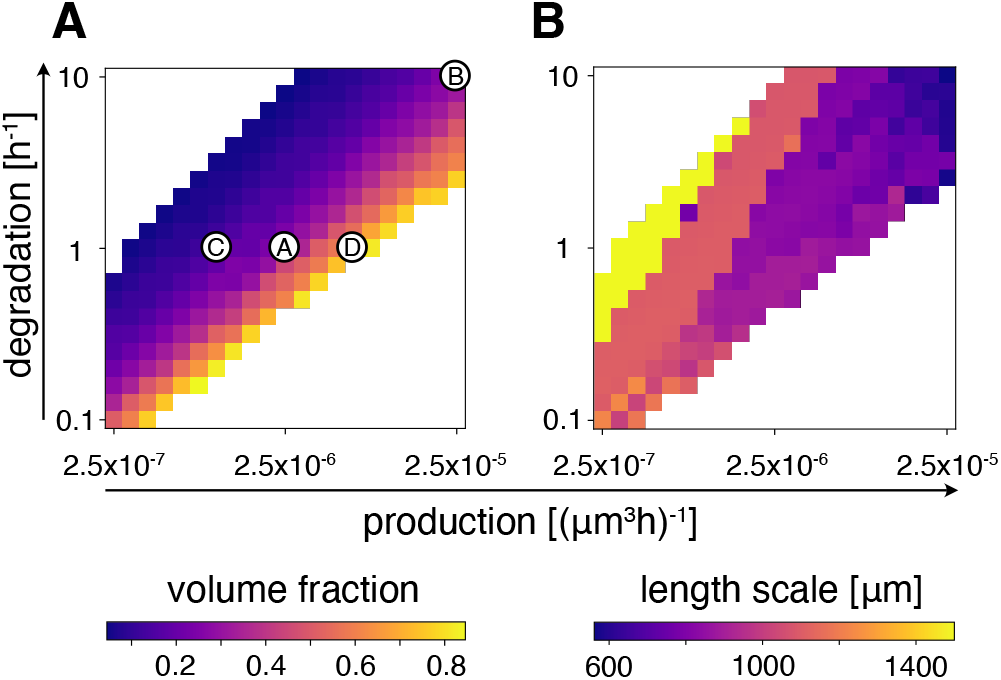
Medulla volume fraction and length scale. The production of thymocytes and the degradation of activated thymocytes regulate both the medulla volume fraction and the typical length scale of medulla-cortex patterns. (*A*) The volume fraction of medullary regions (characterized by a mTEC density *ρ*_M_ *> µ*) is larger for systems with higher production and lower degradation rates. (*B*) The typical length scale of the medulla increases together with the time scale of production and degradation. Medulla morphologies displayed in Fig. 3 are marked with corresponding encircled letters (*A-D*). Simulations are performed in two dimensions due to the high computational demand of three-dimensional simulations; *p* and *d* are varied over the indicated parameter range, and the remaining parameters are chosen as shown in Tab I.

In contrast, an increase (decrease) of both the production and degradation rates decreases (increases) the typical length scale of the cortex–medulla architecture (Fig. 5*B*) while keeping the volume fraction (approximately) constant. Thus, we conclude that the timescale of thymocyte production and degradation fixes the length scale of the cortico-medullary pattern. This correlation between pattern length scale and the time scale of the thymocyte production and degradation rate can be explained by the fact that mTECs proliferate only where thymocytes are selected negatively; this is because only activated thymocytes can induce mTEC growth [2, 3]. Hence, the medullary tissue adapts to the supply of thymocytes: While mTEC-driven negative selection is restricted to the interior of the thymus medulla, thymocytes are only supplied from outside the medulla (the cortex). Accordingly, the width of medullary regions is limited because, in wider medullae, the degradation in their interior cannot be balanced by the surface influx^1^. To illustrate the morphological change brought about by the imposed length scale and medullary volume fraction, we mark the specific parameters used in Fig. 3 in Fig. 5. Thus, interestingly, while the production (positive selection) of SP thymocytes and the activation and subsequent degradation (negative selection) of thymocytes is central to thymic function, in our model it also elicits the non-trivial convoluted shape of the mouse thymus medulla and prohibits mTEC accumulation into a single, compact central medulla.

While the time scale of thymocyte production and degradation sets the typical length scale of the cortex–medulla pattern, changes over an order of magnitude are needed to appreciably modulate it (Fig. 5*B*) and are, thus, likely are not accessible experimentally. However, experimentally, varying medullary volume fractions have been achieved using knock-out mice. For instance, *H2-Aa*^*-/-*^ mice, that cannot produce αβ-CD4 SP thymocytes (the majority of thymocytes that give differentiation signals to the medulla [2, 3, 14]), possess significantly reduced medullary volume and the medulla consists solely of medullary islets [7]. It has been speculated that another, rarer, type of thymocyte is co-responsible for stimulating mTEC-proliferation [14]. Thus, this observation agrees with our prediction for low thymocyte numbers (Fig. 3*C*).

Experimental results by Irla et al. [3] show that a higher fraction of self-reactive to self-tolerant thymocytes in the entire SP thymocyte pool leads to larger medullae and mTEC numbers. Assuming that the higher fraction implies higher absolute numbers of self-reactive thymocytes, the increased medullary volume fraction is in line with the model prediction. However, to our knowledge, the effect of ‘titrating’ the concentration of self-reactive thymocytes on medullary morphology has not been reported so far.

Autoreactive thymocytes can be rescued from deletion [59], in particular due to sparse antigen encounter [60], by knocking out the proapoptotic protein Bim. Because activated thymocytes can escape death, this might lengthen the time that they transmit growth signals to their surrounding mTECs. Thus, *Bim*^*-/-*^ mice may have a larger medullary volume fraction under the assumptions of our model. Should this not be the case, the growth signal that activated thymocytes transmit to mTECs may be down-regulated before the thymocytes die under wt conditions. Within our model’s description of the system, degradation would then be the transition out of the signaling state and not synonymous with thymocyte death.

As detailed in section II, DCs and macrophages are responsible for the phagocytosis of activated thymocytes and strongly impact thymocyte death [29–31]. Consequently, modulating the thymocyte degradation strength might be possible via recruiting phagocytes to, or excluding them from, the thymus. In particular, it was shown that increased chemotaxis of phagocytes into the thymus leads to higher phagocyte densities and stronger negative selection [53, 61, 62], while decreasing the chemotaxis of phagocytes into the thymus can decrease phagocyte densities [63]. However to the authors’ knowledge, there are no quantitative analyses of the medulla morphology in such mouse models.

In summary, the model predicts that the production of self-reactive thymocytes and their degradation in the activated state set the medullary volume fraction and induce a characteristic length scale of the cortex-medulla pattern. This, in turn, means that by altering the production and degradation of thymocytes the medullary morphology can be changed in experiments.

### D. The effect of medullary structure on negative selection

We have discussed how cortex and medulla segregate and how the convoluted cortex–medulla structures form. Finally, we ask whether this segregation—based on our spatial thymic cross-talk model—could have a functional advantage for thymocyte selection. According to Irla et al. [7], the convoluted morphology potentially facilitates the rapid deletion of thymocytes. Based on our theoretical analysis, we hypothesize that the stimulation of mTEC growth by thymocytes not only regulates the overall ratio of mTECs to thymocytes but also promotes the self-organization of a spatial structure within the medulla that enhances the efficiency of negative selection.

Enhancing the rate of negative selection—that is, the rate of recognition of autoreactive thymocytes—reduces the likelihood of these cells escaping into the body before their deletion because fewer self-reactive cells remain that can escape. Concurrently, the thymic medulla exhibits a considerable turnover of mTECs, which is reported to be as high as 13% per day [39, 64]. Consequently, the maintenance of this tissue demands high metabolic resources, which underscores the necessity for efficient tissue utilization [65]. As a proxy for negative selection, we define the effective recognition rate of self-reactive thymocytes as the volume-averaged rate of recognition in the thymus of a certain morphology

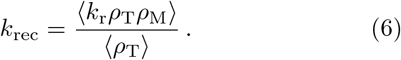

To assess the impact of medullary morphology on the recognition of thymocytes, we test different fixed medulla structures (switching off medullary dynamics) with the same total amount of medullary tissue as in the simulation with the full self-organizing dynamics (Fig. 6*A*).

**Figure 6.**
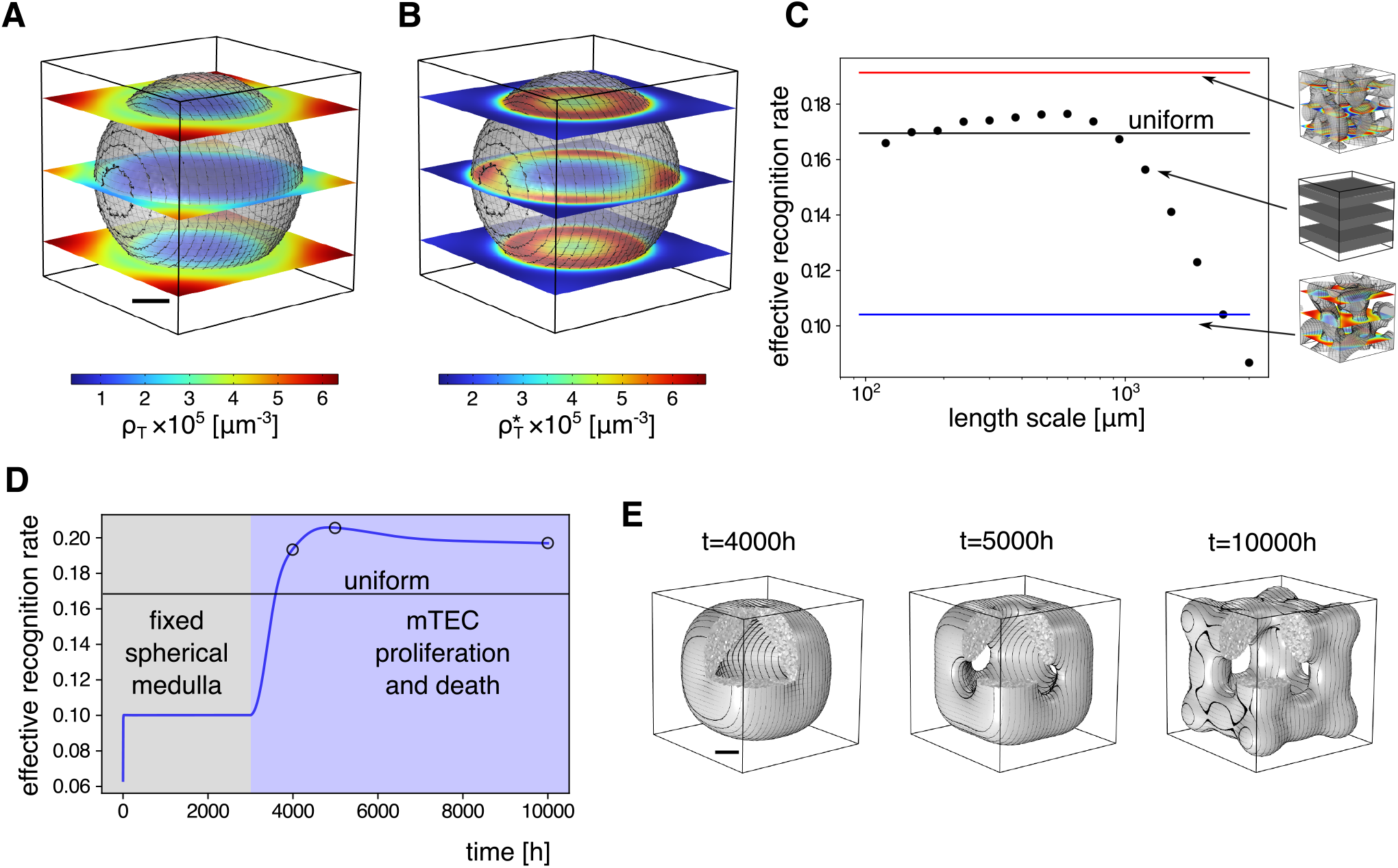
Medullary structure impacts the efficiency of negative selection. With a single spherical medulla, *ρ*_T_ accumulates mainly outside the medullary region (*A*), and 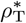 is located almost exclusively at the edge of the medullary region (*B*), regions with *ρ*_M_ *>* 10^−4^µm^−3^ shaded in gray. (*C*) The effective recognition rate *k*_rec_ of medullary lamellae with different length scales (dots; inset) are compared with the effective recognition rate in a homogeneous thymus with the same amount of mTECs but without medulla–cortex segregation (black), and self-organized medullae with (red) and without (blue) chemotaxis. (*D*) The effective thymocyte recognition rate for a single sphere (gray-shaded region) is below that of a homogeneous thymus, while it quickly rises above this threshold once the mTEC growth- and death-dynamics are turned on (blue-shaded region). Blue: *k*_rec_, as calculated from simulation, black: *k*_rec_, as calculated from homogeneous densities of the average density ⟨*ρ*_M_⟩. (*E*) Snap-shots are shown for the time points labeled in (*D*), where regions with *ρ*_M_ *> µ* are colored in gray. The scale bar length is 500µm.

As the simplest possible form, we consider a spherical medulla placed in the simulated volume (Fig. 6)*A, B*. The effective recognition rate in this scenario (*k*_rec_ ≈ 0.1h^−1^, Fig. 6*D* for *t <* 3000h) is lower than in the case in which all cell densities are uniformly distributed, that is, without any segregation of cortical and medullary tissue (*k*_rec_ ≈ 0.17h^−1^ Fig. 6*C*, green line). This is due to the compact form of the medulla increasing the mean distance from the cortical region into the medulla. This increased distance results in longer times for the thymocytes to reach the medulla [7] and, consequently, an accumulation of thymocytes in the cortex (Fig. 6*A*). Moreover, thymocytes are mostly recognized and degraded at the edge of the medulla (Fig. 6*B*), and thus, a large portion of medullary volume does not participate in negative selection. Both aspects show that the characteristic length scale of the cortico–medullary pattern influences the effective recognition rate. The effective deletion rate in the wild-type simulation is larger than the rate without cortex–medulla segregation (*k*_rec_ ≈ 0.19h^−1^, Fig. 6*C*, red line).

This simulation shows that cortex–medulla segregation can be favorable when compared to an unstructured thymus, but only if the morphology assembles at the right length scale. To systematically analyze the impact of different length scales on the effective recognition rate, we study the effective recognition rate for lamellar medullae alternating with lamellae of cortex at a given length scale.

For large length scales, the effective recognition rate is low, as in the case of the spherical medulla. On the other hand, for very short length scales, the rate approaches the value for a well-mixed system (Fig. 6*C*, blue dots). At intermediate length scales roughly comparable with the length scale of the thymocyte gradients, the rate surpasses the effective recognition rate for the well-mixed system.

The heuristic argument for this behavior is that for small length scales of the lamellae (around 200 µm) the chemokine and thymocyte gradients are weak compared to the cortex–medulla length scale. This averages out the cortex–medulla segregation resulting in an effective recognition rate close to the value without any segregation (Fig. 6*C*, black dots). For increasing length scales, chemotaxis is strongest (c.f. SI Appendix Sec. IV.B) and co-localizes the thymocytes with high mTEC densitiesin-creasing the effective deletion rate. At even larger length scales ≳ 1 mm, the effective recognition rate decreases again as for the single spherical medulla. Importantly, without chemotaxis inducing co-localization, even the simulation including medulla self-organization (Fig. 6*C*, blue line) gives a lower effective recognition rate than the well-mixed system (Fig. 6*C*, black line). This finding agrees well with experimental results on strong autoimmunity in *Ccr7*^*-/-*^ and CCR7 ligand-deficient *plt* mice caused by a lack of medullary accumulation of thymocytes [66].

Given that length scales which are much shorter or longer than the wt pattern length scale reduce the effective recognition rate, it is interesting to ask how the self-organization process based on thymic cross-talk ensures appropriate, intermediate length scales. Due to the thymocyte-transmitted proliferation signal, the medullary tissue adapts to coincide with high densities of activated thymocytes. Thymic cross-talk thus ensures that medullary tissue only forms where it is reached by thymocytes (c.f. Sec. III C) and no “superfluous” regions of thymic medulla (as in Fig. 6*C*) are created. To illustrate, we initialize the system under wild-type parameters with a single, fixed spherical medulla before later activating the proliferation signal (mTEC growth and death dynamics) (Fig. 6 (*D,E*)). This rapidly reorganizes the medulla shape. The length scale from the cortex into the medulla decreases and the effective recognition rate increases close to the value reached in the wild-type simulation.

Thymocyte production and the deletion of autoreactive thymocytes are central thymic functions. Coupling the thymocyte-selection process to mTEC proliferation in our model, allows for the formation of a spatial medulla structure that increases the selection efficiency by thymocyte colocalization due to chemotaxis.

Experiments on the spatial distribution of the different cell populations can test the proposed dynamics. For instance, as illustrated by the example of the spherical medulla (Fig. 6*A, B*), a thymus with medullary regions that have been widened artificially through a process unconnected to the negative selection of the observed self-reactive population would show a decrease in self-reactive cell densities in the interior of the medulla.

## IV. CONCLUSIONS

During ontogeny, the spatial structure of organs has to organize by itself driven by cellular interactions, the mechanics of tissue growth, and guiding cues. In the murine thymus, the discovery of thymic cross-talk [5, 6] showed that the intricate spatial structure of medulla and cortex depends on signaling between the thymic tissue and motile thymocytes migrating through the tissue. Diverse genetic perturbations affect this spatial organization [15]. Here, we introduce a spatially explicit model based on the basic experimentally verified interactions within thymic cross-talk. This theoretical model demonstrates how these interactions can lead to the self-organization of a convoluted medulla–cortex structure, as illustrated in Fig. 3.

This spatial thymic cross-talk model predicts that the division of the thymus into cortex and medulla is caused by the combined effect of a chemotactic aggregation instability, as known from Keller–Segel models [34], and a positive feedback in the mTEC proliferation dynamics leading to a growth-induced instability [41]. While chemotaxis localizes thymocytes to the medulla, without chemotaxis thymocytes accumulate in the cortex (Fig. 4); this finding recapitulates experimental results in mice with suppressed CCR7–mediated chemotaxis [18, 26, 28, 50]. The depletion of thymocytes in the medulla strongly suggests that the formation of the medulla in these knockouts is not based on chemotaxis via another signaling axis. Rather, our model suggests a chemotaxis-independent Turing-like pattern formation mechanism, driven by the mTEC proliferation dynamics.

The resulting medulla–cortex pattern is characterized by the width of alternating medullary and cortical regions and the volume fraction of the medulla (Figs. 3, 5). The growth of medullary tissue depends on a thymocyte-dependent proliferation signal. Therefore, medullae form in regions where thymocytes are present. Thus, the medulla shape and the width of medullary islets and branches adapt to the supply of thymocytes. As a result, the intrinsic pattern length scale depends upon the production and degradation of thymocytes, a phenomenon known from Keller–Segel models [67], as well as phase-separating and reaction–diffusion systems with production–degradation dynamics [58, 68]. In addition, the total amount of thymocytes sets the volume fraction occupied by the medulla. Due to the combination of these effects, the model recapitulates the three-dimensional convoluted structure of wild-type medullae [7, 8] and the insular medullary structure found in *H2-Aa*^*-/-*^ mice [7].

As our model describes the changes in volume fraction purely in terms of mTEC number changes, a model extension is necessary to describe morphological changes in which the total mTEC numbers do not change, such as in *Ltα*^*-/-*^ mice [7, 53]. We expect that experimental measurements of changes in local cell densities of mTEC and thymocyte subpopulations inside the medulla will elucidate these additional effects.

Because the spatial cross-talk model describes the self-adaptation of the medulla to the thymocyte supply, we expect that the concepts discussed here will help understand such additional processes as well as the role of spatial heterogeneity within the cortex and medulla, see e.g., Ref. [7]. Furthermore, we expect that the model can be extended to analyze the effects of different initial conditions [42], tissue-growth mechanics, aging of the thymus, see e.g., Ref. [69], and the interplay of more finely resolved mTEC and thymocyte species [4].

Does the self-organized structure affect the function of the thymus, i.e., the thymocyte selection process? Our analysis shows that, in the model, the average deletion rate of autoreactive thymocytes is morphology dependent: Considering different medullary morphologies and comparing them with a structureless organ, we find an efficient medulla–cortex structure that allows for a high effective deletion rate of autoreactive thymocytes (Fig. 6). The robust formation of this efficient structure is ensured by coupling the spatial self-organization to the selection process: The thymocyte-dependent mTEC proliferation signal leads to medulla growth in those regions where thymocytes have to be selected. Thus, function elicits form, and in turn, form also follows function. It remains an interesting open question whether the spatial architecture affects not only the average rate of selection but also the T cell receptor repertoire formation.

Medullary cross-talk forms a feedback control for the selection process, because activated thymocytes are a proxy for the supply of self-reactive thymocytes that have to be recognized in the medulla. Our model suggests that in the thymus, this control not only regulates the relative numbers of the interacting cell types [11] but also their spatial arrangement. Exciting future directions for research include extended theoretical modeling to resolve intra-medullary structure and experimental work probing the suggested morphogenetic process, as well as possible model extensions. In particular, the self-organization of cortex–medulla patterns “from scratch” in thymic organoids (reaggregate thymic organ cultures) will be fascinating to study in detail. Moreover, it is an intriguing question whether other organs rely on similar direct feedback to robustly self-organize functional spatial architectures.

## ACKNOWLEDGMENTS

We thank Heather Melichar and Marilaine Fournier, Fridtjof Brauns, Laeschkir Wuärthner, Andriy Goychuck, and Kieran James for fruitful discussions. We acknowledge financial support by the Deutsche Forschungsgemeinschaft (DFG, German Research Foundation) through the Excellence Cluster ORIGINS under Germany’s Excellence Strategy (EXC-2094-390783311), and the Chan-Zuckerberg Initiative (CZI). This research was conducted within the Max-Planck’School ‘Matter to Life’ supported by the German Federal Ministry of Education and Research (BMBF) in collaboration with the Max Planck Society.

## Footnotes

1 The process discussed here is more broadly known as droplet or plateau splitting and selects, together with interrupted coars- ening at shorter length scales, a range of stable pattern length scales [58], further detailed in an upcoming publication [**?** ].

