## Supplementary Information for "Basic interactions responsible for thymus function explain convoluted medulla shape"

### Supplementary Information
### Basic interactions responsible for thymic function explain convoluted medulla shape


David Muramatsu,[1,*] Henrik Weyer,[1,*] Florian Gartner,[1] and Erwin Frey[1,2,†]

[1]*Arnold Sommerfeld Center for Theoretical Physics and Center for NanoScience, Department of Physics, Ludwig-Maximilians-Universität München, Theresienstraße 37, D-80333 München, Germany*

[2]*Max Planck School Matter to Life, Hofgartenstraße 8, D-80539 Munich, Germany*



(Dated: August 9, 2024)


#### I. EXPERIMENTAL BASIS FOR THE MODEL AND PARAMETER CHOICE

In the main text, we proposed a coarse-grained model of the most important interactions of thymic cross-talk that take part in medulla formation. In this section, we expand on the experimental evidence of the interactions and model parameters outlined in the main text. First, we discuss the simplifications of the known interactions and constituents of the thymus, focusing on the interactions taking place within the medulla in Sec. I A. Subsequently, in Sec. I B, we explain the parameter choices used in the simulations based on the experimental literature. Finally, in Sec. I C, we derive the sensitivity function that modulates the chemotaxis in dependence of chemokine densities in the medulla, following an argument by Stevens and Othmer [1].

##### A. Model Abstractions

###### 1. *Chemokines*

Both cTECs and mTECs, as well as dendritic cells, emit chemokines that steer thymocytes in various stages into their appropriate microenvironments (for an overview of these interactions, see [2, 3]). In this publication, we mainly model interactions between autoreactive SP thymocytes and medullary tissue. Thus, we do not expect the production of chemokines from cTECs which act on earlier development stadia of thymocytes to play a role in these processes. Rather, these effects are subsumed in the effective, cortex-restricted production term of thymocytes (the term $p\phi$ in Eq. (1) in the main text).

SP and/or post-positive selection DP thymocytes are attracted to mTEC-produced chemokines CCL21 [4], CCL19 [5], CCL1 [6], CCL8 [6], EBI2-ligand 7α25-OHC [7], as well as the mTEC- and DC-produced chemokines CCL17 and CCL22 [8–10]. While many of these chemokines aid in recruiting thymocytes to the medulla [6, 7, 9, 11–15], only CCL21 has so far been shown to significantly impact medullary morphology and size [6–9, 11, 12, 16, 17]. At the same time, DCs are attracted to the medulla via diverse chemokines, see e.g. [18–21], with the DC subsets that express the highest levels of CCL17 and CCL22 also expressing CCR7, the chemokine receptor for CCL21 [10] and cDCs, and the most numerous DC subset being mainly located in the medulla [18, 22, 23].

Thus, as all these attractive chemokine gradients predominantly originate in the medulla, we conflate them into one generalized chemokine gradient.

###### 2. *mTECs*

At a coarse level, mTECs can be divided into the $\text{mTEC}^{\text{lo}}$ and $\text{mTEC}^{\text{hi}}$ populations which express the two surface markers MHC-II and CD80 at low and high levels, respectively [24]. These two populations can be split up into further phenotypes [10, 25–29] whose differentiation and proliferation is controlled by interaction with thymocytes [30–33].

These cell populations are functionally distinct and recent evidence points to two branches of differentiation within the mTEC population originating from the same progenitor [29]. Within the proposed model, one branch differentiates into an $\text{Aire}^+$ cell line [29]. As $\text{Aire}^+$ cells as a whole are presenting nearly all of the bodies protein [26, 34], this branch is tasked with inducing self-tolerance [35]. $\text{Aire}^+$ cells are known to be denser at the CMJ, particularly in young mice, and this preference is attenuated in adult mice [32]. The effect of this distribution on thymocyte localization, as well as the mechanism leading to this distribution of $\text{Aire}^+$ cells is a fascinating extant question that we do not cover here.

The other branch is marked by a high expression of CCL21 and Cytokeratin 5 (Krt5) [29]. While a type of cell expressing high amounts of CCL21 has been identified to be enriched at the CMJ [36], Krt5 is used as a medullary marker [37], with $\text{Krt5}^+$ cells evenly distributed in the medulla, and only known to be abnormally distributed in clusters in *Fezf2*$^{-/-}$ mice [38]. Thus, the production of the chemokine by a single mTEC density nearly homogeneous within the medullary region in the model appears reasonable.

---


* These authors contributed equally.
†

##### 3. Antigen presenting cells

Apart from mTECs, several discrete cell subsets are known to present antigen to thymocytes for negative selection in the medulla. While mTECs, DCs and B-Cells [34, 39] clonally delete thymocytes, only mTECs are known to induce a stimulatory capacity for the mTEC-proliferation signal in thymocytes [31]. As we model activated thymocytes to possess stimulatory capacity for mTEC proliferation, only mTEC-activated thymocytes fall into this subset, while B cell and DC-activated thymocytes do not.

The fraction of DC-stimulated cells is estimated to make up between 30 and 70% of negatively selected thymocytes [40–44]. For simplicity, in this publication, we assume the value to be 50%. As the vast majority of negative selection events in the medulla are split between DCs and mTECs [41] in two-photon microscopy experiments, we neglect the effect of B cells and only consider mTECs and DCs.

DC-activation of thymocytes (and the minor contribution due to B cells) can be seen as an effective degradation term of thymocytes that leads to an effective reduction of the production rate of thymocytes, see also Sec. I B 2, I B 5. Specifically, because DCs mostly localize to the medulla, see e.g. Ref. [18, 45], we approximate their density $\rho_{\mathrm{DC}}$ as being proportional to the mTEC density, i.e., one has $\rho_{\mathrm{DC}} \sim \rho_{\mathrm{M}}$. Therefore, thymocyte deletion by DCs can be modeled by $-k_{\mathrm{DC}}\rho_{\mathrm{T}}\rho_{\mathrm{DC}} \approx -\tilde{k}_{\mathrm{DC}}\rho_{\mathrm{T}}\rho_{\mathrm{M}}$. Given the rate of thymocyte recognition by mTECs $\bar{k}_{\mathrm{r}}$ and that mTEC recognition accounts roughly for one half of negatively selected thymocytes, one has $\bar{k}_{\mathrm{r}} \approx \tilde{k}_{\mathrm{DC}}$. Because only mTEC-activated thymocytes stimulate mTEC growth, only these thymocytes are converted into the activated model species $\rho^*_{\mathrm{T}}$. As a result, the model equations for the thymocytes that include DC-mediated negative selection, can be approximated as

$$\partial_t \rho_{\mathrm{T}} = D_{\mathrm{T}}\nabla^2\rho_{\mathrm{T}} - \nabla(\rho_{\mathrm{T}}\mathbf{v}_{\mathrm{C}}) - 2\bar{k}_{\mathrm{r}}\rho_{\mathrm{T}}\rho_{\mathrm{M}} + \bar{p}\phi(\rho_{\mathrm{M}})\,, \quad (1)$$

$$\partial_t \rho^*_{\mathrm{T}} = D_{\mathrm{T}}\nabla^2\rho^*_{\mathrm{T}} - \nabla(\rho^*_{\mathrm{T}}\mathbf{v}_{\mathrm{C}}) + \bar{k}_{\mathrm{r}}\rho_{\mathrm{T}}\rho_{\mathrm{M}} - d\,\rho^*_{\mathrm{T}}\,. \quad (2)$$

Here, the thymocytes are produced with an average cortical production rate $\bar{p}$. Rescacaling the thymocyte density as $\tilde{\rho}_{\mathrm{T}} = \frac{\rho_{\mathrm{T}}}{2}$ leads to an effectively halved production rate, and a doubled conversion rate $2\bar{k}_{\mathrm{r}}\rho_{\mathrm{M}}$ which is the same for autoreactive and activated thymocytes:

$$\partial_t \tilde{\rho}_{\mathrm{T}} = D_{\mathrm{T}}\nabla^2\tilde{\rho}_{\mathrm{T}} - \nabla(\tilde{\rho}_{\mathrm{T}}\mathbf{v}_{\mathrm{C}}) - 2\bar{k}_{\mathrm{r}}\tilde{\rho}_{\mathrm{T}}\rho_{\mathrm{M}} + \frac{\bar{p}}{2}\phi(\rho_{\mathrm{M}})\,, \quad (3)$$

$$\partial_t \rho^*_{\mathrm{T}} = D_{\mathrm{T}}\nabla^2\rho^*_{\mathrm{T}} - \nabla(\rho^*_{\mathrm{T}}\mathbf{v}_{\mathrm{C}}) + 2\bar{k}_{\mathrm{r}}\tilde{\rho}_{\mathrm{T}}\rho_{\mathrm{M}} - d\,\rho^*_{\mathrm{T}}\,. \quad (4)$$

Thus, re-writing $\tilde{\rho}_{\mathrm{T}} \to \rho_{\mathrm{T}}$, one arrives at Eq. (1, 2) of the main text with $k_{\mathrm{r}} = 2\bar{k}_{\mathrm{r}}$ and $p = \frac{\bar{p}}{2}$.

##### 4. Thymocytes

T cells developing in the thymus consist of several subsets, out of which the $\alpha\beta - \mathrm{CD4}^+$ and $\alpha\beta - \mathrm{CD8}^+$ single positive (SP) subsets are the most abundant [39]. Out of these, CD4 SP thymocytes possess the dominant stimulatory capacity for mTEC growth [31, 32], with other subsets playing a subordinate role [46]. In this publication, we mainly focus on the functional role of thymocytes for mTEC differentiation, and thus, model all thymocytes that induce mTEC proliferation upon antigen recognition as the same effective cell species.

When considering the spatial distribution of thymocytes with respect to their environment, we introduce a population of tolerant thymocytes, as they are not degraded due to negative selection, and thus have different kinetics. For simplicity, we use the parameters of CD4 SP tolerant thymocytes for this population. However, all tolerant thymocytes have the same underlying kinetics of production in the cortex, transition to the medulla, and eventual exit out of the thymus, and thus share the same phenomenology.

CD4 SP thymocytes that are self-reactive also include the subset of regulatory T cells ($\mathrm{T_{reg}}$). Since, to our knowledge, there exists no data on the stimulatory capacity of $\mathrm{T_{reg}}$ cells on mTEC proliferation and differentiation, we do not explicitly model this subset. However, $\mathrm{T_{reg}}$ cells are known to preferentially arise in response to recognizing sparsely presented antigen [47], such that their kinetics may be different from general negatively selected CD4 SP thymocytes. Thus, if $\mathrm{T_{reg}}$ cells do possess stimulatory capacity in thymic cross-talk, an additional thymocyte population with stimulatory capacity might need to be introduced, potentially yielding interesting new phenomenology regarding the respective cell density distributions.

#### B. Parameters

The model parameters used for the results displayed in the main text are based on experimental findings, as we detail below. Taken together, the rates for the simulation reflecting the wild-type conditions can be found in Table I of the main text and for all other simulations in Tables S1, S2 of the SI.

##### 1. Medullary and cortical volume

The volume of one thymic lobe has been estimated at 25 - 40mm$^3$ [48, 49], whereas the medullary volume have been measured to be around 10% (2.5mm$^3$) [48] to 25% (10mm$^3$) [49]. Thus, the cortical volume ranges between 90% (23mm$^3$) [48] and 75% (30mm$^3$) [49]. Although the thymic volume itself is not a parameter in the analysis performed in this publication, it is necessary to estimate the rate of autoreactive thymocyte production.

##### 2. Production of autoreactive thymocytes

The production rate of autoreactive thymocytes has not been quantified. We therefore estimate it from measured selection rates by assuming a steady-state turnover: The rates for the selection of autoreactive thymocytes in the DP and SP stadium have been estimated experimentally in Refs. [50, 51], in good accordance with the fraction of autoreactive DP and SP thymocytes that undergo negative selection measured by Ref. [40]. Since in our model, the inflow of autoreactive thymocytes is equal to the outflow by negative selection in steady state, the average rate of selection of one thymocyte differentiation stage must be equal to the average rate of production of thymocytes that are selected in this stadium.

Commonly, only CD4 SP thymocytes are thought to impact medullary growth, see e.g. Refs. [31, 40, 51]. Thus, we consider only this thymocyte population to stimulate mTEC growth in our model. Stritesky et al. [51] found the total rate of negative selection — and thus the total production if we assume a steady-state turnover of the thymocytes — of autoreactive CD4 SP thymocytes to be $p_{\text{tot}} = 1.66 \cdot 10^5 \text{h}^{-1}$. With a cortical volume $V_{\text{C}} = 23 - 30\text{mm}^3$, the lower bound for the production rate of self-reactive thymocytes is

$$\bar{p} = \frac{p_{\text{tot}}}{V_{\text{C}}} = \frac{1.66 \cdot 10^5}{30 \cdot 10^9} \text{h}^{-1}\,\mu\text{m}^{-3} \approx 5 \cdot 10^{-6} \text{h}^{-1}\,\mu\text{m}^{-3}\,. \quad (5)$$

As discussed in Sec. I A 3, we estimate the total number of mTEC-stimulating activated thymocytes to be half that of total activated CD4 SP thymocytes, as the other half is activated by DCs. Thus, the production rate of thymocytes that undergo selection by mTECs, is

$$p = \frac{\bar{p}}{2} = 2.5 \cdot 10^{-6} \text{h}^{-1}\,\mu\text{m}^{-3}\,. \quad (6)$$

We note that post-positive selection DP thymcoytes can localize into the medulla [9, 15]. However, to our knowledge, it is not conclusively known whether these cells can drive mTEC differentiation. Thus, depending on the extent of both localization and differentiation signaling by post-positive selection DP thymocytes, the thymocyte production rate may have to be corrected to a larger value.

##### 3. Thymocyte production in the cortex

DP thymocytes are located in the cortex, where they receive signals for positive selection [2]. After receiving these signals, they differentiate and transition into the medulla [9, 11, 13–15]. To reflect this dynamic in our model, we localize SP thymocyte production to the cortex using the function

$$\phi(\mathbf{x}) = \frac{1}{1 + (\rho_M(\mathbf{x})/\mu)^2}\,, \quad (7)$$

with the parameter $\mu$ chosen such that the function falls off from one to zero around $\rho_{\text{M}} \sim \mu$. We discuss the choice of $\mu$ in Sec. I B 12.

##### 4. Production rate of tolerant CD4 SP thymocytes

CD4 SP thymocytes survive negative selection at approximately half the rate with which they are negatively selected [51]. However, since we are only considering half of the negatively selected thymocytes in their production rate $p$, we approximate the production rate of tolerant thymocytes as $p_{\text{T}}^0 = p$.

##### 5. Antigen recognition

In the thymus medulla, antigen is presented at different frequencies, with some antigens being expressed ubiquitously, and others, so-called *tissue restricted antigens*, are expressed sparsely [34]. The timing and extent of the negative selection against ubiquitous (OVAp) [52] and sparse antigen (RIPmOVA) [53] have been analyzed in overlay experiments, where specific thymocytes are overlaid onto thymic slices containing their cognate antigen. In the case of ubiquitous antigen, the activation of thymocytes is almost immediate, with a two–hour lag-period, after which the phagocytosis by randomly encountered phagocytes sets in [52]. In the experiment with sparse antigen selection, the lag extends beyond three hours, but in both experiments, deletion happens within the first nine hours, and there is little change in living cell numbers afterward [52, 53].

We choose our rates to be roughly in accordance with these results, with the mTEC-based activation rate at $\bar{k}_{\text{r}} = \frac{1}{2}\text{h}^{-1}$, and thus, the effective activation rate

$$k_{\text{r}} = 2\,\bar{k}_{\text{r}} = 1\,\text{h}^{-1}\,, \quad (8)$$

as detailed in Sec. I A 3.

The validity of the RIPmOVA system as a model for sparse antigen presentation has been challenged [47, 54, 55], and lower rates of antigen presentation than the ones that are prevalent in this system would lead to lower rates of antigen recognition.

To estimate a lower bound for the thymic selection rate, we consider the fraction of autoreactive T cells in the entire body. While it has been suggested that around one-third of the body's T cell repertoire is autoreactive [47, 56, 57], it is likely that almost all thymocytes that are autoreactive against ubiquitous antigen and those which react strongly against sparse antigen are selected in the thymus [47, 56]. Thus, we assume the time scale of selection against these thymocytes to be at least an order of magnitude faster than the time scale of thymocyte egress (around 4-5 days [58]), as otherwise, a large fraction of these autoreactive thymocytes would escape.

Nonetheless, it is conceivable that the rate of thymocyte selection is within the span of 1/1h to 1/5d, the rate of thymic egress. The fundamental mechanism underlying our model is not abolished by this parameter and a convoluted medulla forms due to thymocyte production and degradation on both ends of the rate's spectrum.

##### 6. *Decay rate of activated thymocytes*

Upon antigen recognition, thymocytes upregulate phagocytosis-inducing factors [52], and thymocyte counts decrease sharply after the appearance of the first death markers, and from the data presented in [52], we estimate a half-life of activated thymocytes of one hour.

##### 7. *Exit rate of SP thymocytes*

SP thymocytes exit the thymus with a rate of approximately five days [58], if they are not negatively selected. Thymocytes can exit the thymus after maturing from both cortex and medulla [11, 59]. Thus, for simplicity we choose a uniform exit rate $k_{\text{ex}} = (5\text{d})^{-1} \approx 0.008\text{h}^{-1}$.

##### 8. *mTEC death rate*

The half-life of mTECs has been estimated between 0.9 w for young mice, and 6 - 11 w for adult mice [60–62], yielding a decay rate of $k_{\text{m}} = 0.0004 - 0.005\text{h}^{-1}$. Both ends of the thus given spectrum yield qualitatively the same morphology, as long as the ratio of mTEC proliferation rate $k_{\text{b}}$ and mTEC death rate $k_{\text{m}}$ are held constant. We choose the faster dynamics with a value of $k_{\text{m}} = 0.005\text{h}^{-1}$, representing the mTEC dynamics in young mice [62] as the patterning of the thymus will mostly happen during this stage.

##### 9. *Medullary mTEC density*

The density of mTECs in the medulla has been measured at $2 - 3 \cdot 10^{-4} \mu\text{m}^{-3}$ for 5wk and around $2.5 \cdot 10^{-4} \mu\text{m}^{-3}$ for 12wk old mice by Ref. [63] and around $8 \cdot 10^{-4} \mu\text{m}^{-1}$ by Ref. [64] at ten days after birth, the former of which are in good agreement with values by Ref. [49] who found a total number of mTECs of $\sim 2.5 \cdot 10^6$ cells and a thymic right lobe volume of approximately $10\text{mm}^3$, 5wk after birth. We thus adopt this value of $\rho_{\text{M}} = 2.5 \cdot 10^{-4} \mu\text{m}^{-3}$ in our model parameters.

##### 10. *mTEC proliferation rate*

The mTEC proliferation rate $k_{\text{b}} = 3 \cdot 10^3 \mu\text{m}^3\,\text{h}^{-1}$ is chosen to reproduce a medullary mTEC density of $\rho_{\text{M}} = 2.5 \cdot 10^{-4} \mu\text{m}^{-3}$ in the wt steady state (see Sec. I B 9), with a three-fold higher medullary density allowed by the carrying capacity.

##### 11. *mTEC carrying capacity*

The carrying capacity $K$ limits the production of new mTECs and thus sets the density above which mTECs do not proliferate. In early embryonic development, the mTEC densities are about three-fold higher [64] than at the stage of development in which the three-dimensional structure of the thymus was analyzed [48, 65], implying a considerable range of possible densities. We chose $K = 7.5 \cdot 10^{-4} \mu\text{m}^{-3}$ such that mTEC densities are capped but a three-fold increase in mTEC densities compared to the chosen wild-type density $2.5 \cdot 10^{-4} \mu\text{m}^{-3}$ is allowed.

##### 12. *mTEC density scale*

As mentioned in Sec. I B 9, we choose the mTEC density to be $\rho_{\text{M}} \approx 2.5 \cdot 10^{-4} \mu\text{m}^{-3}$ inside the medulla. Thus, we choose a value of $\mu = 10^{-4} \mu\text{m}^{-3}$ for the mTEC density scale.

##### 13. *Chemokine decay rate*

The rate of chemokine decay due to proteases has been estimated between $0.72\text{h}^{-1}$ and $18\text{h}^{-1}$ by Ref. [66] based on previous studies [67, 68]. However, since chemokines may be taken up by their cognate receptors and atypical chemokine receptors [69], this rate may be larger. Here, we use a rough estimate of $k_{\text{e}} = 10\text{h}^{-1}$. While ten-fold larger values impose narrower cortical regions, we checked that they do not qualitatively change the medullary morphology in the wild-type case.

##### 14. *Chemokine production rate*

The average concentration of CCL21 has been measured to be $\sim$ 5ng per mg thymus by Ref. [70] and 2ng/mg by Ref. [71]. At a mass of CCL21 of 14.5 kDa [72], this translates to densities of 160−400nM. Using the chemokine decay rate, as well as the medullary mTEC densities described above, we choose our chemokine production rate to result in the approximate average density of about 200nM in the wild-type steady-state and arrive at a production rate of $1.25 \cdot 10^{-2}\text{M} \cdot \mu\text{m}^3\text{h}^{-1}$.

##### 15. *Chemokine Diffusivity*

Chemokine diffusivity in serum has been estimated at $6.5 \cdot 10^{-7}\text{cm}^2\text{s}{-1} = 2.34 \cdot 10^5 \mu\text{m}^2\text{h}^{-1}$ in Ref. [66] based on experimental data from Ref. [73–75]. Since the chemokine diffuses through the interstitium, we use a ten-fold lower value, however, choosing the original value does not change the morphology of the system for the chosen 'wt' parameters.

###### 16. Thymocyte Diffusivity

The thymocyte motility has been measured to be approximately $D_{\mathrm{T}} = 400\mu\mathrm{m}^2\mathrm{min}^{-1} = 24 \cdot 10^3 \mu\mathrm{m}^2\mathrm{h}^{-1}$ by Ref. [76], which we use in our numerical analysis.

Tolerant thymocytes that do not undergo chemotaxis are distributed approximately evenly in cortex and medulla in our model, even though they are produced in the cortex. This can be explained as follows: Tolerant thymocytes spend a time of $t_{\mathrm{med}} = 4 - 5\mathrm{d}$ inside the thymus medulla [58], see Sec. I B 7. As without chemotaxis, these cells perform random motion with the above given diffusivity, within that time they cover a distance of about $l_{\mathrm{diff}} \sim \sqrt{t_{\mathrm{med}} \cdot D_{\mathrm{T}}} = \sqrt{5 \cdot 24\mathrm{h} \cdot 24 \cdot 10^3 \mu\mathrm{m}^2\mathrm{h}^{-1}} \approx 1.7\mathrm{mm}$, or about half the chosen simulated volume. Thus, a time of five days is enough for equilibrating the tolerant thymocyte density between the cortex, where they are produced, and the medulla.

###### 17. Chemotaxis strength

The strength of chemotaxis $T$ is a free parameter, since we model the combined effect of the chemokines, nor is it known for chemotaxis in the thymic environment. In the simulations, we choose a value of $10^5\mu\mathrm{m}^2\mathrm{h}^{-1}$ which yields the desired patterns.

###### 18. Chemokine binding affinity

The equilibrium constant of receptor-chemokine binding for CCL21-CCR7 has been estimated between 1.6 and 10nM [77–79], here we take the choice of 5nM, orienting ourselves on the estimate by Ref. [66], which we use as a generic chemokine binding constant.

##### C. Chemotaxis sensitivity function

Chemokine receptor binding kinetics have previously been successfully used to model CCR7-dependent T cell and DC chemotaxis [66, 80]. We, also include this type of signaling as a chemokine density-dependent sensitivity function $\chi(\rho_{\mathrm{C}})$ in the advection term of our model.

Therefore, we choose simple receptor binding kinetics [81] of the form

$$R + C \underset{k_{\mathrm{off}}}{\overset{k_{\mathrm{on}}}{\rightleftharpoons}} R \cdot C\,, \tag{9}$$

where $R$ and $C$ signify unbound receptor and chemokine molecules, and $R{\cdot}C$ signifies a bound receptor-chemokine complex, following the derivation by Stevens and Othmer [1]. In steady state, binding and unbinding are balanced, which reads following mass-action kinetics

$$k_{\mathrm{on}}[R][C] = k_{\mathrm{on}}\left([R_{\mathrm{tot}}] - [R \cdot C]\right)[C] = k_{\mathrm{off}}[R \cdot C]\,, \tag{10}$$

such that

$$[R \cdot C] = \frac{k_{\mathrm{on}}\,[R_{\mathrm{tot}}]\,[C]}{k_{\mathrm{off}} + k_{\mathrm{on}}[C]} = [R_{\mathrm{tot}}]\,\frac{[C]}{K_{\mathrm{OffOn}} + [C]}\,, \tag{11}$$

where $K_{\mathrm{OffOn}} = {}^{k_{\mathrm{off}}}/_{k_{\mathrm{on}}}$ is the dissociation constant of chemokine-receptor binding and $[R_{\mathrm{tot}}]$ is the concentration of all bound and unbound receptors in the cell we consider. Since the cell senses its surrounding chemokine gradient via the gradient of bound receptors on its surface, we calculate

$$\begin{aligned}\nabla[R \cdot C] &= \nabla\,[R_{\mathrm{tot}}]\,\frac{[C]}{K_{\mathrm{OffOn}} + [C]}\\ &= [R_{\mathrm{tot}}]\,\frac{K_{\mathrm{OffOn}}}{\left(K_{\mathrm{OffOn}} + [C]\right)^2}\nabla[C]\,,\end{aligned} \tag{12}$$

such that the total receptor density $[R_{\mathrm{tot}}]$ can be absorbed into the factor for chemotaxis strength, $T$, and the sensitivity function depends only on the chemokine density $[C] = \rho_C$,

$$\chi(\rho_C) = \frac{K_{\mathrm{OffOn}}}{\left(K_{\mathrm{OffOn}} + \rho_C\right)^2}\,. \tag{13}$$

#### II. LINEAR STABILITY ANALYSIS AND MODEL REDUCTION

In this section, we present the mathematical analysis of the model equations. In the first subsection, we discuss the linear stability analysis (LSA) of the full model. The LSA allows one to determine the parameter region in which uniform density distributions of the cells and the chemoattractant (the homogeneous steady state, HSS) become unstable due to spatially inhomogenenous perturbations (lateral instability). In this regime, the system spontaneously segregates into cortical and medullary regions (cf. gray-shaded region in main text Fig. 4($A$)).

In the following two subsections, we reduce the full model to a chemotaxis-free system and to a system without Turing instability. Using quasi–steady–state (QSS) approximations, we relate these systems to two-component Keller–Segel and Turing reaction–diffusion systems. This reduction shows explicitly that the two instability mechanisms of the full model, discussed in Sec. III.A in the main text, are the chemotaxis-driven instability conceptualized by the Keller–Segel model and the Turing reaction–diffusion instability.

##### A. Original model

In our spatial thymic cross-talk model, the densities of thymocytes, activated thymocytes, mTECs, and chemokines are given by the equations [cf. Eqs. (1-4) in

| parameter name | symbol | wt values | Fig. 3 | | | | Fig. 4 | | unit |
|---|---|---|---|---|---|---|---|---|---|
| | | | $(A)$ | $(B)$ | $(C)$ | $(D)$ | with chemotaxis | without chemotaxis | |
| chemokine diffusivity | $D_{\mathrm{C}}$ | 23400 | | | | | | | $\mu\mathrm{m}^2\,\mathrm{h}^{-1}$ |
| thymocyte random motility | $D_{\mathrm{T}}$ | 24000 | | | | | | | $\mu\mathrm{m}^2\,\mathrm{h}^{-1}$ |
| taxis strength | $T$ | $10^5$ | | | | | | 0 | $\mu\mathrm{m}^2\,\mathrm{h}^{-1}$ |
| mTEC random motility | $D_{\mathrm{M}}$ | 3 | | | | | | | $\mu\mathrm{m}^2\,\mathrm{h}^{-1}$ |
| receptor binding constant | $K_{\mathrm{OffOn}}$ | 5 | | | | | | | nM |
| thymocyte production | $p$ | $2.5 \cdot 10^{-6}$ | | $2.5 \cdot 10^{-5}$ | $10^{-6}$ | $7 \cdot 10^{-6}$ | | | $\mu\mathrm{m}^{-3}\,\mathrm{h}^{-1}$ |
| mTEC density scale | $\mu$ | $10^{-4}$ | | | | | | | $\mu\mathrm{m}^{-3}$ |
| thymocyte deletion | $d$ | 1 | | 10 | | | | | $\mathrm{h}^{-1}$ |
| thymocyte conversion | $k_{\mathrm{r}}$ | $1.5 \cdot 10^3$ | | | | | | | $\mu\mathrm{m}^3\,\mathrm{h}^{-1}$ |
| mTEC birth rate | $k_{\mathrm{b}}$ | $3 \cdot 10^3$ | | | | | | | $\mu\mathrm{m}^3\,\mathrm{h}^{-1}$ |
| mTEC carrying capacity | $K$ | $7.5 \cdot 10^{-4}$ | | | | | | | $\mu\mathrm{m}^{-3}$ |
| mTEC decay | $k_{\mathrm{m}}$ | 0.005 | | | | | | | $\mathrm{h}^{-1}$ |
| chemokine emission | $k_{\mathrm{e}}$ | $1.4 \cdot 10^{-2}$ | | | | | | | $\mathrm{M}\,\mu\mathrm{m}^3\,\mathrm{h}^{-1}$ |
| chemokine decay | $k_{\mathrm{c}}$ | 10 | | | | | | | $\mathrm{h}^{-1}$ |
| tolerant thymocyte production | $p_{\mathrm{T}}^0$ | - | - | - | - | - | $2.5 \cdot 10^{-6}$ | $2.5 \cdot 10^{-6}$ | $\mu\mathrm{m}^{-3}\,\mathrm{h}^{-1}$ |
| thymocyte exit | $k_{\mathrm{ex}}$ | - | - | - | - | - | 1/5 | 1/5 | $\mathrm{d}^{-1}$ |
| system side length | $L$ | 3 | | | | | | | mm |
| simulated time | $t_{\mathrm{max}}$ | 2000 | | | | | | | h |

Table S1. Parameters for wt (third column; cf. Fig. 3($A$) in the main text), and for Fig. 3, and Fig. 4 from the main text. For Figs. 3, 4, the parameters varying from the wt values are shown. Parameters that are not part of the specific model variant are indicated with a dash.

| parameter name | symbol | wt values | Fig. 5 | Fig. 6 | | unit |
|---|---|---|---|---|---|---|
| | | | | sphere – ($A$, $B$, $D$, $E$) | lamellae – ($C$) | |
| chemokine diffusivity | $D_{\mathrm{C}}$ | 23400 | | | | $\mu\mathrm{m}^2\,\mathrm{h}^{-1}$ |
| thymocyte random motility | $D_{\mathrm{T}}$ | 24000 | | | | $\mu\mathrm{m}^2\,\mathrm{h}^{-1}$ |
| taxis strength | $T$ | $10^5$ | | | | $\mu\mathrm{m}^2\,\mathrm{h}^{-1}$ |
| mTEC random motility | $D_{\mathrm{M}}$ | 3 | | | | $\mu\mathrm{m}^2\,\mathrm{h}^{-1}$ |
| receptor binding constant | $K_{\mathrm{OffOn}}$ | 5 | | | | nM |
| thymocyte production | $p$ | $2.5 \cdot 10^{-6}$ | $2.5 \cdot 10^{-7} - 2.5 \cdot 10^{-6}$ | | | $\mu\mathrm{m}^{-3}\,\mathrm{h}^{-1}$ |
| mTEC density scale | $\mu$ | $10^{-4}$ | | | | $\mu\mathrm{m}^{-3}$ |
| thymocyte deletion | $d$ | 1 | $0.1 - 10$ | | | $\mathrm{h}^{-1}$ |
| thymocyte conversion | $k_{\mathrm{r}}$ | $1.5 \cdot 10^3$ | | | | $\mu\mathrm{m}^3\,\mathrm{h}^{-1}$ |
| mTEC birth rate | $k_{\mathrm{b}}$ | $3 \cdot 10^3$ | | | | $\mu\mathrm{m}^3\,\mathrm{h}^{-1}$ |
| mTEC carrying capacity | $K$ | $7.5 \cdot 10^{-4}$ | | | | $\mu\mathrm{m}^{-3}$ |
| mTEC decay | $k_{\mathrm{m}}$ | 0.005 | | | | $\mathrm{h}^{-1}$ |
| chemokine emission | $k_{\mathrm{e}}$ | $1.4 \cdot 10^{-2}$ | | | | $\mathrm{M}\,\mu\mathrm{m}^3\,\mathrm{h}^{-1}$ |
| chemokine decay | $k_{\mathrm{c}}$ | 10 | | | | $\mathrm{h}^{-1}$ |
| pattern length scale | $L_{\mathrm{p}}$ | - | - | - | $0.12 - 3$ | mm |
| equilibration time | $T_{\mathrm{eq}}$ | - | - | 3000 | - | h |
| system side length | $L$ | 3 | | | 1 | mm |
| simulated time | $t_{\mathrm{max}}$ | 2000 | | $10^4$ | | h |

Table S2. Parameters for wt (third column; cf. Fig. 3($A$) in the main text), and for Fig. 5, and Fig. 6 from the main text. Parameters varying from the wt values are shown. Parameters that are not part of the specific model variant are indicated with a dash.

the main text]

$$\partial_t \rho_{\mathrm{T}} = D_{\mathrm{T}} \nabla^2 \rho_{\mathrm{T}} - \nabla(\rho_{\mathrm{T}} v_{\mathrm{C}}) - k_{\mathrm{r}} \rho_{\mathrm{T}} \rho_{\mathrm{M}} + p\, \phi(\rho_{\mathrm{M}}) \,, \tag{14a}$$

$$\partial_t \rho_{\mathrm{T}}^* = D_{\mathrm{T}} \nabla^2 \rho_{\mathrm{T}}^* - \nabla(\rho_{\mathrm{T}}^* v_{\mathrm{C}}) + k_{\mathrm{r}} \rho_{\mathrm{T}} \rho_{\mathrm{M}} - d\, \rho_{\mathrm{T}}^* \,, \tag{14b}$$

$$\partial_t \rho_{\mathrm{M}} = D_{\mathrm{M}} \nabla^2 \rho_{\mathrm{M}} + k_{\mathrm{b}} \rho_{\mathrm{T}}^* \rho_{\mathrm{M}} \Big(1 - \frac{\rho_{\mathrm{M}}}{K}\Big) - k_{\mathrm{m}} \rho_{\mathrm{M}} \,, \tag{14c}$$

$$\partial_t \rho_{\mathrm{C}} = D_{\mathrm{C}} \nabla^2 \rho_{\mathrm{C}} + k_{\mathrm{e}} \rho_{\mathrm{M}} - k_{\mathrm{c}} \rho_{\mathrm{C}} \,, \tag{14d}$$

where $\phi(\rho_{\mathrm{M}}) = (1 + (\rho_{\mathrm{M}}/\mu)^2)^{-1}$ and $\mathbf{v}_{\mathrm{C}} = T\chi(\rho_{\mathrm{C}})\nabla\rho_{\mathrm{C}}$ have been introduced in the main text. Defining the vector $\boldsymbol{\rho} = (\rho_{\mathrm{T}}, \rho_{\mathrm{T}}^*, \rho_{\mathrm{M}}, \rho_{\mathrm{C}})^T$, these equations can be written more compactly as

$$\partial_t \boldsymbol{\rho} = \nabla \left[\mathbf{M}(\boldsymbol{\rho}) \nabla \boldsymbol{\rho}\right] + \mathbf{f}(\boldsymbol{\rho}) \,. \tag{15}$$

Here, the mobility matrix $\mathbf{M}$ accounts for the diffusion coefficients and chemotaxis terms and $\mathbf{f}$ contains the growth, conversion, and decay terms. Thus, they are given by

$$\mathbf{M}(\boldsymbol{\rho}) = \begin{pmatrix} D_\mathrm{T} & 0 & 0 & -T\chi(\rho_\mathrm{C})\rho_\mathrm{T} \\ 0 & D_\mathrm{T} & 0 & -T\chi(\rho_\mathrm{C})\rho_\mathrm{T}^* \\ 0 & 0 & D_\mathrm{M} & 0 \\ 0 & 0 & 0 & D_\mathrm{C} \end{pmatrix} , \tag{16}$$

$$\mathbf{f}(\boldsymbol{\rho}) = \begin{pmatrix} p\,\phi(\rho_\mathrm{M}) - k_\mathrm{r}\rho_\mathrm{T}\rho_\mathrm{M} \\ k_\mathrm{r}\rho_\mathrm{T}\rho_\mathrm{M} - d\,\rho_\mathrm{T}^* \\ k_\mathrm{b}\rho_\mathrm{T}^*\rho_\mathrm{M}\,(1-\rho_\mathrm{M}/K) - k_\mathrm{m}\rho_\mathrm{M} \\ k_\mathrm{e}\rho_\mathrm{M} - k_\mathrm{c}\rho_\mathrm{C} \end{pmatrix} . \tag{17}$$

To understand in which parameter regimes the model spontaneously segregates cortex and medulla, we perform a linear stability analysis of the homogeneous steady state (HSS) using Wolfram Mathematica. The general procedure is as follows:

1. Determine the homogeneous steady state (HSS) that represents the stationary configuration of the system with uniform cell and chemoattractant densities. The HSS with positive densities for all components is given for the wild-type parameters (Tab. I in the main text) by $\hat{\rho}_\mathrm{T} = 3.94 \cdot 10^{-5}\mu\mathrm{m}^{-3}$, $\hat{\rho}_\mathrm{T}^* = 3.63 \cdot 10^{-6}\mu\mathrm{m}^{-3}$, $\hat{\rho}_\mathrm{M} = 6.14 \cdot 10^{-5}\mu\mathrm{m}^{-3}$, $\hat{\rho}_\mathrm{C} = 85.98\mathrm{nM}$.

2. Linearize Eq. (15) around this HSS for small perturbations $\delta\rho_i = \rho_i(\mathbf{x},t) - \hat{\rho}_i$, $i = \mathrm{T}, \mathrm{T}^*, \mathrm{M}, \mathrm{C}$.

3. Solve the linearized system using a normal mode decomposition. This yields the growth rates $\sigma_q^i$, $i = 1,2,3,4$ of the normal modes with mode number $q$. These indicate which perturbations around the HSS grow or decay (see below).

The HSS densities $\boldsymbol{\rho}_\mathrm{hss}$ (given above for the wild-type parameters) are found by solving $0 = \mathbf{f}(\boldsymbol{\rho}_\mathrm{hss})$ because all gradients vanish in the case of uniform densities. The linearized dynamics are obtained by inserting $\boldsymbol{\rho} = \boldsymbol{\rho}_\mathrm{hss} + \delta\boldsymbol{\rho}$ in Eq. (15) and keeping only those terms linear in $\delta\boldsymbol{\rho}$. This yields

$$\partial_t \delta\boldsymbol{\rho} = \left[\mathbf{M}(\boldsymbol{\rho}_\mathrm{hss})\nabla^2 + \mathbf{J}\right] \delta\boldsymbol{\rho} , \tag{18}$$

with the Jacobian

$$J_{ij} = \partial_{\rho_j} f_i(\boldsymbol{\rho})\big|_{\boldsymbol{\rho}=\boldsymbol{\rho}_\mathrm{hss}} . \tag{19}$$

The linearized dynamics, Eq. 18, together with the periodic boundary conditions, can then be solved using a normal mode decomposition (Fourier series)

$$\delta\boldsymbol{\rho}(\mathbf{x},t) = \sum_\mathbf{q} e^{\sigma_q t} e^{i\mathbf{q}\mathbf{x}} \delta\boldsymbol{\rho}_\mathbf{q} . \tag{20}$$

Here, the growth rate $\sigma_q$ for a Fourier mode with wavenumber $q = |\mathbf{q}|$ solves the eigenvalue equation

$$\sigma_q \delta\boldsymbol{\rho}_\mathbf{q} = \left[-q^2\mathbf{M}(\boldsymbol{\rho}_\mathrm{hss}) + \mathbf{J}\right] \delta\boldsymbol{\rho}_\mathbf{q} . \tag{21}$$

This eigenvalue equation for the growth rates $\sigma_q$ can be solved employing Wolfram Mathematica's `Eigenvalue[]` function for the matrix $\left[-q^2\mathbf{M}(\boldsymbol{\rho}_\mathrm{hss}) + \mathbf{J}\right]$. Solving this eigenvalue equation, one obtains four growth rates $\sigma_q^i$ with $i = 1,2,3,4$ because the diagonalized matrix is four-dimensional. The real part of the eigenvalues indicates whether the amplitude of the corresponding perturbation mode grows ($\mathrm{Re}[\sigma_q^i] < 0$) or decays ($\mathrm{Re}[\sigma_q^i] > 0$) [cf. Eq. (20)]. To analyze the stability of the HSS, one defines the dispersion relation $\sigma_q$ as the growth rate with the maximal real part for a given wavenumber $q$.

For parameter values for which the real part of the growth rates is negative for all wavenumbers $q$, $\mathrm{Re}[\sigma_q] < 0$, the HSS is linearly stable. In contrast, in the case that $\mathrm{Re}[\sigma_q] > 0$ holds for a band of wavenumbers $q \in (q_-, q_+)$ with $q_+ > q_- \geq 0$, small deviations from the HSS with wavenumbers $q \in (q_-, q_+)$ will grow over time and the HSS is unstable. Because the HSS is only unstable due to the growth of spatially inhomogenenous perturbations, this instability is called a lateral instability. For the analyzed parameter region, one finds $q_- > 0$ because the HSS is locally stable, i.e., $\mathrm{Re}[\sigma_0] < 0$. Consequently, the lateral instability is of type I within the classification scheme of Cross and Hohenberg [82]. Figure 4($A$) in the main text shows the resulting parameter regime in which the system is laterally unstable (gray-shaded region).

We now discuss the two mechanisms underlying this lateral instability in the spatial cross-talk model. To this end, we study two reduced models that either exclude chemotaxis or the reaction–diffusion instability.

##### B. Chemotaxis-free model

Figure 4($A$–$B$) of the main text shows that the spatial cross-talk model predicts that the thymus tissue segregates into cortex and medulla even without chemotaxis. To analyze the lateral instability in this case [blue-shaded parameter region in Fig. 4($A$)], we set the chemotaxis strength $T$ in Eqs. (14) to zero which yields the chemotaxis-free system

$$\partial_t \rho_\mathrm{T}(\mathbf{x},t) = D_\mathrm{T}\nabla^2\rho_\mathrm{T} - k_\mathrm{r}\rho_\mathrm{T}\rho_\mathrm{M} + p\,\phi(\rho_\mathrm{M}) , \tag{22a}$$

$$\partial_t \rho_\mathrm{T}^*(\mathbf{x},t) = D_\mathrm{T}\nabla^2\rho_\mathrm{T}^* + k_\mathrm{r}\rho_\mathrm{T}\rho_\mathrm{M} - d\,\rho_\mathrm{T}^* , \tag{22b}$$

$$\partial_t \rho_\mathrm{M}(\mathbf{x},t) = D_M\nabla^2\rho_\mathrm{M} + k_\mathrm{b}\rho_\mathrm{T}^*\rho_\mathrm{M}\left(1 - \frac{\rho_\mathrm{M}}{K}\right) - k_\mathrm{m}\rho_\mathrm{M} . \tag{22c}$$

This model contains a positive feedback loop based on the growth and decay of mTECs (cf. Sec. III.A in the main text), which we show explicitly to induce a reaction–diffusion (Turing) instability in this section. As discussed in the main text, the feedback loop is due to the increased production of activated thymocytes in regions of increased mTEC densities by recognition (first reaction term in Eq. (22b)). In return, the activated thymocytes stimulate further proliferation of the mTECs, enhancing initial variations in the mTEC density (first reaction term in Eq. (22c)).

To simplify the feedback mediated by the production of activated thymocytes, we will neglect their time evolution. We assume that the density of activated thymocytes adapts instantaneously to changes in the densities of the mTECs and the non-activated thymocytes. Thus, we approximate the density $\rho_\mathrm{T}^*$ by its quasi-steady-state (QSS) distribution $\tilde{\rho}_\mathrm{T}^*$ obtained by setting $\partial_t \tilde{\rho}_\mathrm{T}^* = 0$ in Eq. (22b). While this approximation does not give the exact temporal dynamics of the system, it does not change the steady-state patterns.

To calculate the QSS density distribution, we define the linear operator $\mathcal{L}^{-1} \equiv (1 - D_\mathrm{T}/d\,\nabla^2)^{-1}$.[1] Then, starting from Eq. (22b) and setting $\partial_t \tilde{\rho}_\mathrm{T}^* = 0$, one obtains the QSS distribution of the activated thymocytes $\tilde{\rho}_\mathrm{T}^*$ as

$$\tilde{\rho}_\mathrm{T}^* = \frac{k_\mathrm{r}}{d}\mathcal{L}^{-1}\rho_\mathrm{T}\rho_\mathrm{M}. \tag{23}$$

Note that the operator $\mathcal{L}^{-1}$ is the identity when acting on uniform densities. Therefore, to separate its uniform contribution from the one depending on gradients in the density fields $\rho_\mathrm{T,M}$, we rewrite the operator as

$$\mathcal{L}^{-1} \equiv 1 + \frac{D_\mathrm{T}}{d}\mathcal{D}. \tag{24}$$

Here, the linear operator $\mathcal{D}$ defined by this equation can be expanded for weak gradients as $\mathcal{D} = \nabla^2 + \mathcal{O}(\nabla^4)$. Thus, $\mathcal{D}$ can be understood as a generalized diffusion operator that approaches the Laplacian $\nabla^2$ in the long-wavelength limit. Inserting the quasi-steady state distribution Eq. (23) into the mTEC dynamics Eq. (22c), one obtaines the reduced system

$$\partial_t\rho_\mathrm{T}(\mathbf{x},t) = D_\mathrm{T}\nabla^2\rho_\mathrm{T} - k_\mathrm{r}\rho_\mathrm{T}\rho_\mathrm{M} + p\,\phi(\rho_\mathrm{M})\,, \tag{25a}$$

$$\begin{aligned}\partial_t\rho_\mathrm{M}(\mathbf{x},t) = {}& D_\mathrm{M}\nabla^2\rho_\mathrm{M} + \frac{D_\mathrm{T}k_\mathrm{b}k_\mathrm{r}}{d^2}\rho_\mathrm{M}\Big(1-\frac{\rho_\mathrm{M}}{K}\Big)\mathcal{D}\rho_\mathrm{T}\rho_\mathrm{M} \\ &+ \frac{k_\mathrm{b}k_\mathrm{r}}{d}\rho_\mathrm{T}\rho_\mathrm{M}^2\Big(1-\frac{\rho_\mathrm{M}}{K}\Big) - k_\mathrm{m}\rho_\mathrm{M}\,.\end{aligned} \tag{25b}$$

Because the thymocytes diffuse much more quickly than the sedentary mTECs, gradients in the mTEC density are large compared to gradients in the thymocyte density $\rho_\mathrm{T}$, allowing one to approximate $\mathcal{D}\rho_\mathrm{T}\rho_\mathrm{M} \approx \rho_\mathrm{T}\mathcal{D}\rho_\mathrm{M}$. Using this approximation, we define the effective mTEC diffusion coefficient

$$D_\mathrm{M}^\mathrm{eff}(\rho_\mathrm{T},\rho_\mathrm{M}) \equiv \frac{D_\mathrm{T}k_\mathrm{b}k_\mathrm{r}}{d^2}\rho_\mathrm{T}\rho_\mathrm{M}\Big(1-\frac{\rho_\mathrm{M}}{K}\Big)\,. \tag{26}$$

Linearizing around the homogeneous steady state $\boldsymbol{\rho}_\mathrm{hss}$, which is unaffected by the approximations performed in this section, the diffusion coefficient $D_\mathrm{M}^\mathrm{eff}(\rho_\mathrm{T},\rho_\mathrm{M})$ is set constant to its value $D_\mathrm{M}^\mathrm{eff} \equiv D_\mathrm{M}^\mathrm{eff}(\rho_\mathrm{T,hss},\rho_\mathrm{M,hss})$ evaluated at the densities $\boldsymbol{\rho}_\mathrm{hss}$. We then arrive at the generalized two-component reaction–diffusion system

$$\partial_t\rho_\mathrm{T}(\mathbf{x},t) = D_\mathrm{T}\nabla^2\rho_\mathrm{T} - k_\mathrm{r}\rho_\mathrm{T}\rho_\mathrm{M} + p\,\phi(\rho_\mathrm{M})\,, \tag{27a}$$

$$\begin{aligned}\partial_t\rho_\mathrm{M}(\mathbf{x},t) = {}& \big[D_\mathrm{M}\nabla^2 + D_\mathrm{M}^\mathrm{eff}\mathcal{D}\big]\,\rho_\mathrm{M} \\ &+ \frac{k_\mathrm{b}k_\mathrm{r}}{d}\rho_\mathrm{T}\rho_\mathrm{M}^2\Big(1-\frac{\rho_\mathrm{M}}{K}\Big) - k_\mathrm{m}\rho_\mathrm{M}\,.\end{aligned} \tag{27b}$$

In this system, the thymocytes (density $\rho_\mathrm{T}$) are produced and degraded based on the mTEC density $\rho_\mathrm{M}$. In turn, the mTEC density decays linearly due to cell death ($\sim k_\mathrm{m}$) but has a T-cell dependent second-order self-production term $\sim \rho_\mathrm{T}\rho_\mathrm{M}^2$ which stems from the feedback loop, i.e., the local part of $\tilde{\rho}_\mathrm{T}^*$ [cf. Eqs. (23), (24)]. The non-local part of $\tilde{\rho}_\mathrm{T}^*$, i.e., the diffusion of the activated thymocytes, reduces to the additional diffusion term with the effective diffusion coefficient $D_\mathrm{M}^\mathrm{eff}$ for the mTECs. While the reactions give rise to a stable fixed point $\mathrm{Re}[\sigma_0] < 0$, the system is laterally unstable due to a Turing instability [83]. This Turing instability is possible because the thymocytes diffuse much more quickly than the mTECs as the effective diffusion constant $D_\mathrm{M}^\mathrm{eff}$ is about two orders of magnitude smaller than $D_\mathrm{T}$ ($D_\mathrm{M}$ is even smaller). While the two-component system Eqs. (27) is not an exact reaction–diffusion system due to the generalized diffusion operator $\mathcal{D}$, in the long-wavelength limit, one can use $\mathcal{D} \approx \nabla^2$ to arrive at

$$\partial_t\rho_\mathrm{T}(\mathbf{x},t) = D_\mathrm{T}\nabla^2\rho_\mathrm{T} - k_\mathrm{r}\rho_\mathrm{T}\rho_\mathrm{M} + p\,\phi(\rho_\mathrm{M})\,, \tag{28a}$$

$$\begin{aligned}\partial_t\rho_\mathrm{M}(\mathbf{x},t) = {}& \big(D_\mathrm{M} + D_\mathrm{M}^\mathrm{eff}\big)\,\nabla^2\rho_\mathrm{M} \\ &+ \frac{k_\mathrm{b}k_\mathrm{r}}{d}\rho_\mathrm{T}\rho_\mathrm{M}^2\Big(1-\frac{\rho_\mathrm{M}}{K}\Big) - k_\mathrm{m}\rho_\mathrm{M}\,.\end{aligned} \tag{28b}$$

Figure S1 shows the dispersion relations $\sigma_q$ obtained for the wild-type parameters with the full chemotaxis-free model Eq. (22), the reduced model with generalized diffusion Eq. (27), and the complete reduction to the Turing system Eq. (28). All systems show a band of unstable modes. The QSS approximation (and $\mathcal{D}\rho_\mathrm{T}\rho_\mathrm{M} \approx \rho_\mathrm{T}\mathcal{D}\rho_\mathrm{M}$) only introduces small deviations in the dispersion relation. The reduced reaction–diffusion system Eqs. (28) approximates the dispersion relation of the full system well at long wavelengths (small wavenumbers $q$) while it deviates strongly at shorter wavelength due to neglecting higher-order gradient terms in the approximation $\mathcal{D} \approx \nabla^2$. Importantly, the reduced system Eqs. (28) retains the emergence of a band of unstable modes $q$ with $\sigma_q > 0$ above a minimal wavenumber $q_-$ while the system is locally stable ($\sigma_0 < 0$). This verifies that the lateral instability of the full chemotaxis-free model is a generalization of the instability found in the reduced two-component system.

Two-component reaction–diffusion systems show a Turing instability if the Segel–Jackson criterion is fulfilled [84]. It requires that the components of the Jacobian of the reaction terms at the hss have a particular combination of signs. The slow-diffusing species must have a positive, the fast-diffusing species a negative diagonal entry. Moreover, the off-diagonal terms must have opposite

[1] The linear operator $\mathcal{L}^{-1}$ acts as the convolution with a Yukawa kernel. Given a function $g(\mathbf{x})$ for $\mathbf{x} \in \mathbb{R}^3$, one has $\mathcal{L}^{-1}g = \int \mathrm{d}^3y K(|\mathbf{x}-\mathbf{y}|)g(\mathbf{y})$ with $K(r) = \exp(-r/\sqrt{D_\mathrm{T}/d})/r$.

signs. The linearization of Eq. (28) gives the Jacobian

$$\mathbf{J}_{\mathrm{reac}} = \begin{pmatrix} -k_{\mathrm{r}}\rho_{\mathrm{M}} & -k_{\mathrm{r}}\rho_{\mathrm{T}} + p\partial_{\rho_{\mathrm{M}}}\phi \\ \frac{k_{\mathrm{b}}k_{\mathrm{r}}}{d}\rho_{\mathrm{M}}^2\left(1-\frac{\rho_{\mathrm{M}}}{K}\right) & 2\frac{k_{\mathrm{b}}k_{\mathrm{r}}}{d}\rho_{\mathrm{T}}\rho_{\mathrm{M}}\left(1-\frac{\rho_{\mathrm{M}}}{K}\right) - k_{\mathrm{m}} \end{pmatrix} \tag{29}$$

with $\partial_{\rho_{\mathrm{M}}}\phi < 0$ and $\rho_{\mathrm{M}} < K$. Thus, the signs of the components are

$$\begin{pmatrix} - & - \\ + & +/- \end{pmatrix}. \tag{30}$$

The second diagonal term becomes positive if the feedback term $\sim \rho_{\mathrm{T}}\rho_{\mathrm{M}}^2$ becomes dominant. Then, the system can undergo a Turing instability. The resulting combination of signs in the Jacobian is called an activator–substrate system or depletion model [85, 86].

Taken together, the reduced two-component reaction–diffusion system Eqs. (28) shows that the feedback mediated by activated thymocytes allows for a Turing instability in the system. This instability drives the segregation of cortex and medulla in the spatial thymic cross-talk model if the thymocytes do not undergo chemotaxis.

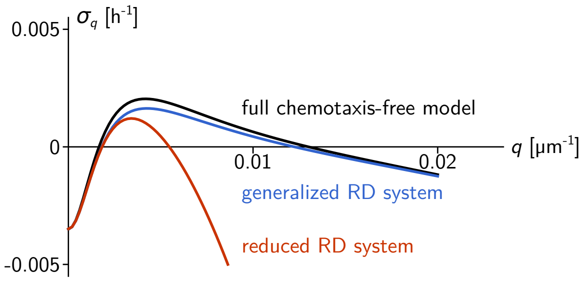


Figure S1. Comparison of the dispersion relations of the chemotaxis-free model Eq. (22) (black), the reduced generalized reaction–diffusion system Eq. (27) (blue), and the reduced reaction–diffusion system Eq. (28) obtained in the long-wavelength limit (red). The dispersion relations are calculated for the wild-type parameters given in Table I.

*Instability mechanism for* $D_{\mathrm{T}} \gg D_{\mathrm{M}}^{\mathrm{eff}}$ — For the wild-type parameters, one has $D_{\mathrm{T}} \gg D_{\mathrm{M}} + D_{\mathrm{M}}^{\mathrm{eff}}$ (by two orders of magnitude). In this case, one can simplify the linear stability analysis by taking the limit $(D_{\mathrm{M}} + D_{\mathrm{M}}^{\mathrm{eff}})/D_{\mathrm{T}} \to 0$. Then, due to their fast diffusion, the thymocyte density $\rho_{\mathrm{T}}$ can be assumed as uniform on the length scales on which the mTEC density varies. Assuming the thymocyte density as constant, the linearized mTEC dynamics Eq. (28b) read

$$\partial_t \delta\rho_{\mathrm{M}}(\mathbf{x},t) = \left(D_{\mathrm{M}} + D_{\mathrm{M}}^{\mathrm{eff}}\right)\nabla^2\delta\rho_{\mathrm{M}} + \left(\partial_{\rho_{\mathrm{M}}}\tilde{f}_{\mathrm{M}}(\boldsymbol{\rho}_{\mathrm{hss}})\right)\delta\rho_{\mathrm{M}}, \tag{31}$$

with $\tilde{f}_{\mathrm{M}}(\boldsymbol{\rho}) = (k_{\mathrm{b}}k_{\mathrm{r}}/d)\rho_{\mathrm{T}}\rho_{\mathrm{M}}^2(1-\rho_{\mathrm{M}}/K) - k_{\mathrm{m}}\rho_{\mathrm{M}}$. Thus, the reaction term is the lower diagonal entry of the full Jacobian Eq. (29). The growth rates $\sigma_q$ for $q > 0$ read

$$\sigma_q = \left(\partial_{\rho_{\mathrm{M}}}\tilde{f}_{\mathrm{M}}(\boldsymbol{\rho}_{\mathrm{hss}})\right) - \left(D_{\mathrm{M}} + D_{\mathrm{M}}^{\mathrm{eff}}\right)q^2, \tag{32}$$

and positive growth rates only occur if

$$\partial_{\rho_{\mathrm{M}}}\tilde{f}_{\mathrm{M}}(\boldsymbol{\rho}_{\mathrm{hss}}) > 0\,, \tag{33}$$

i.e., if the lower diagonal term of the Jacobian is positive. This is the same condition as we obtained from the Segel–Jackson criterion.

As $\tilde{f}_{\mathrm{M}}(\boldsymbol{\rho}_{\mathrm{hss}}) = 0$, an instability occurs only if $\tilde{f}_{\mathrm{M}}$ has a positive slope at one of its roots. The steady state of the full system corresponds to the middle root of $\tilde{f}_{\mathrm{M}}$ as a function of $\rho_{\mathrm{M}}$ [see Fig. S2(*A*)]. Indeed, $\tilde{f}_{\mathrm{M}}$ a function of $\rho_{\mathrm{M}}$ has a positive slope at this root because the positive feedback in mTEC proliferation—mTEC growth $\sim \rho_{\mathrm{T}}\rho_{\mathrm{M}}^2$—induces a cubic shape of $\tilde{f}_{\mathrm{M}}$ as a function of $\rho_{\mathrm{M}}$. Perturbations in the mTEC density around the HSS grow because of the feedback induced by the activated thymocytes: At densities $\rho_{\mathrm{M}} < \rho_{\mathrm{M,hss}}$, the decay of mTECs dominates while proliferation dominates due to the feedback at $\rho_{\mathrm{M}} > \rho_{\mathrm{M,hss}}$ (but $\rho_{\mathrm{M,hss}} \lesssim K$).

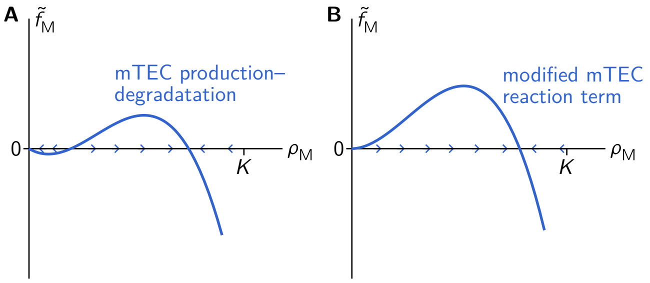


Figure S2. The local mTEC dynamics. (*A*) The production and degradation term $\tilde{f}_{\mathrm{M}}$ of the mTEC density is shown (blue; for constant thymocyte density $\rho_{\mathrm{T}}$). The arrows denote whether the reaction term $\tilde{f}_{\mathrm{M}}$ leads to an increase or decrease of of the mTEC density. (*B*) The modified production and degradation term is shown in which the linear mTEC death term $-k_{\mathrm{m}}\rho_{\mathrm{M}}$ is replaced by the quadratic term $-k_{\mathrm{m}}\rho_{\mathrm{M}}^2/K$.

#### C. Reduced model without Turing instability

In the previous section, we showed that the spatial thymic cross-talk model leads to the segregation into cortex and medulla even without chemotaxis. In this section, we modify the reaction term to exclude this chemotaxis-free (Turing) instability. We then reduce this modified system to a two-component system and show that one obtains Keller–Segel dynamics. Keller–Segel systems also show a lateral instability. Their instability is due to chemotactic aggregation, which shows that the spatial thymic cross-talk model has a Keller–Segel-type chemotactic instability in addition to the Turing instability discussed in the previous section.

To exclude the Turing instability in the reduced two-component reaction–diffusion system Eqs. (28), one has to ensure that the second diagonal term of the Jacobian Eq. (29) is negative at the fixed point. Thus, one has to modify the reaction term $\tilde{f}_{\mathrm{M}}$ such that it does not have a root with positive slope at a density $\rho_{\mathrm{M}} > 0$. In order to not modify the interaction between thymocytes and mTECs, we will only change the mTEC degradation term $k_{\mathrm{m}}\rho_{\mathrm{M}}$. The cubic shape of the reaction term $\tilde{f}_{\mathrm{M}}$ is then lost if we replace the linear mTEC degradation term $-k_{\mathrm{m}}\rho_{\mathrm{M}}$ by the quadratic term $-k_{\mathrm{m}}\rho_{\mathrm{M}}^2/K$ [see Fig. S2($B$)]. Then, mTEC degradation does not dominate at low densities $\rho_{\mathrm{M}}$ while the feedback-induced proliferation dominates at larger densities. Instead, the middle root of $\tilde{f}_{\mathrm{M}}$ is moved to zero.[2]

As a "minimal" modification, we multiply the mTEC decay term with $\rho_{\mathrm{M}}/K$, using the carrying capacity $K$ to nondimensionalize the density. With this modification, we arrive at the modified model without Turing instability (cf. Eqs. (14))

$$\partial_t \rho_{\mathrm{T}} = D_{\mathrm{T}}\nabla^2\rho_{\mathrm{T}} - \nabla(\rho_{\mathrm{T}} v_{\mathrm{C}}) - k_{\mathrm{r}}\rho_{\mathrm{T}}\rho_{\mathrm{M}} + p\,\phi\,, \tag{34a}$$

$$\partial_t \rho_{\mathrm{T}}^* = D_{\mathrm{T}}\nabla^2\rho_{\mathrm{T}}^* - \nabla(\rho_{\mathrm{T}}^* v_{\mathrm{C}}) + k_{\mathrm{r}}\rho_{\mathrm{T}}\rho_{\mathrm{M}} - d\,\rho_{\mathrm{T}}^*\,, \tag{34b}$$

$$\partial_t \rho_{\mathrm{M}} = D_M\nabla^2\rho_{\mathrm{M}} + k_{\mathrm{b}}\rho_{\mathrm{T}}^*\rho_{\mathrm{M}}\Big(1 - \frac{\rho_{\mathrm{M}}}{K}\Big) - k_{\mathrm{m}}\frac{\rho_{\mathrm{M}}^2}{K}\,, \tag{34c}$$

$$\partial_t \rho_{\mathrm{C}} = D_{\mathrm{C}}\nabla^2\rho_{\mathrm{C}} + k_{\mathrm{e}}\rho_{\mathrm{M}} - k_{\mathrm{c}}\rho_{\mathrm{C}}\,. \tag{34d}$$

The linear stability analysis of this model yields the orange-shaded instability region in Fig. 4($A$) in the main text. The system is only laterally unstable if the chemotaxis-strength is finite. This shows that the instability is due to the chemotaxis of the thymocytes.

*Effective Keller–Segel dynamics* — The chemotaxis-driven instability is due to feedback in the chemotactic accumulation of thymocytes: (Activated) thymocytes stimulate mTEC growth. The mTECs produce chemokines that attract more thymocytes to this spatial region. Thus, thymocytes accumulate in this spatial region. The classical Keller–Segel model is the prototypical model showing this chemotaxis-driven instability [87, 88]. Here, we will again use a QSS approximation to reduce the dynamics of the modified model Eqs. (34) to a Keller–Segel model with production and degradation [89].

The Keller–Segel model is a two-component system that models the density of the chemotactic cell density and the density of the chemoattractant, which is produced in dependence on the chemotactic cell density. In the modified model system Eqs. (34), the chemotactic cell density is the total thymocyte density $\rho_{\mathrm{T}}^{\mathrm{tot}} = \rho_{\mathrm{T}} + \rho_{\mathrm{T}}^*$. The thymocytes lead to chemoattractent production indirectly by the stimulation of mTEC growth. To approximate this indirect chemoattractant production by a single chemoattractant production term $p_{\mathrm{C}}(\rho_{\mathrm{T}}^{\mathrm{tot}})$, we determine the local QSS approximations $\tilde{\rho}_{\mathrm{M}}(\rho_{\mathrm{T}}^*)$ and $\tilde{\rho}_{\mathrm{T}}^*(\rho_{\mathrm{T}}^{\mathrm{tot}})$, neglecting the diffusion and chemotaxis of $\rho_{\mathrm{M}}, \rho_{\mathrm{T}}^*$. As the mTECs diffuse very slowly compared to the thymocytes, neglecting their diffusion term will only introduce minor deviations in the chemoattractant production. Neglecting diffusion and chemotaxis of the activated thymocytes $\rho_{\mathrm{T}}^*$ is an uncontrolled approximation, but it will qualitatively capture that, locally, a certain fraction of the thymocytes is activated and those stimulate mTEC proliferation, resulting in an increased chemokine production in regions of high thymocyte density.

The QSS approximation of the mTEC density $\rho_{\mathrm{M}}$ neglecting their diffusion, i.e., setting $D_{\mathrm{M}} = 0$) is obtained by assuming that the mTEC density has fully relaxed to its steady state. As their time evolution vanishes in this (quasi) steady state, it is obtained by setting $\partial_t\rho_{\mathrm{M}} = 0$ in Eq. 34c. This yields[3]

$$\tilde{\rho}_{\mathrm{M}}(\rho_{\mathrm{T}}^*) = K\frac{\rho_{\mathrm{T}}^*}{\frac{k_{\mathrm{m}}}{k_{\mathrm{b}}} + \rho_{\mathrm{T}}^*}\,. \tag{35}$$

Thus, the QSS mTEC density increases with the density of activated thymocytes (as they stimulate mTEC proliferation) up to the carrying capacity $K$. Inserting this expression into Eq. 34b, the local QSS approximation of the activated thymocytes $\rho_{\mathrm{T}}^*$ is obtained analogously by setting $\partial_t\rho_{\mathrm{T}}^* = 0$ (and neglecting the gradient terms). This gives

$$\tilde{\rho}_{\mathrm{T}}^*(\rho_{\mathrm{T}}^{\mathrm{tot}}) = \frac{k_{\mathrm{r}}K}{d + k_{\mathrm{r}}K}\left(\rho_{\mathrm{T}}^{\mathrm{tot}} - \frac{k_{\mathrm{m}}d}{k_{\mathrm{b}}k_{\mathrm{r}}K}\right) \tag{36}$$

if $\rho_{\mathrm{T}}^{\mathrm{tot}} > k_{\mathrm{m}}d/(k_{\mathrm{b}}k_{\mathrm{r}}K)$ and zero otherwise. As a result, the density of activated thymocytes increases linearly with the total thymocyte density if $\rho_{\mathrm{T}}^{\mathrm{tot}} > k_{\mathrm{m}}d/(k_{\mathrm{b}}k_{\mathrm{r}}K)$.

Both relations together yield the monotonously increasing relation $\tilde{\rho}_{\mathrm{M}}(\rho_{\mathrm{T}}^{\mathrm{tot}})$, which results in the two-component Keller–Segel model with thymocyte production and degradation (cf. Ref. [89])

$$\partial_t\rho_{\mathrm{T}}^{\mathrm{tot}} = D_{\mathrm{T}}\nabla^2\rho_{\mathrm{T}}^{\mathrm{tot}} - \nabla(\rho_{\mathrm{T}}^{\mathrm{tot}}v_{\mathrm{C}}) + p\,\tilde{\phi}(\rho_{\mathrm{T}}^{\mathrm{tot}}) - d\,\tilde{\rho}_{\mathrm{T}}^*(\rho_{\mathrm{T}}^{\mathrm{tot}})\,, \tag{37a}$$

$$\partial_t\rho_{\mathrm{C}} = D_{\mathrm{C}}\nabla^2\rho_{\mathrm{C}} + k_{\mathrm{e}}\tilde{\rho}_{\mathrm{M}}(\rho_{\mathrm{T}}^{\mathrm{tot}}) - k_{\mathrm{c}}\rho_{\mathrm{C}}\,, \tag{37b}$$

[2] We assume that the feedback term dominates over the quadratic degradation term. Otherwise, the mTEC density is driven to zero everywhere because one would have $\tilde{f}_{\mathrm{M}} < 0$ for all $\rho_{\mathrm{M}} > 0$ (see Fig. S2($B$)).

[3] Note that this reduction is not possible without the modification of the reaction term that excludes the Turing instability. Without the modification, $\tilde{f}_{\mathrm{M}}$ one root more, which is the middle root with a positive slope that gives rise to the Turing instability. This additional root introduces an effective bistability of the mTEC dynamics because low-density regions of the mTEC pattern relax toward the root $\rho_{\mathrm{M}} = 0$ while high-density regions relax toward the third, high-density root of $\tilde{f}_{\mathrm{M}}$ [see Fig. S2]. The reason is that both the root at $\rho_{\mathrm{M}} = 0$ and the high-density root are stable in the original mTEC dynamics (keeping all other fields fixed). In contrast, only the high-density root is stable in the modified dynamics. Thus, the QSS mTEC density is given by the single (stable) root $\tilde{\rho}_{\mathrm{M}}(\rho_{\mathrm{T}}^*)$. Without the modification of the reaction term, one would have to consider different roots for different regions of the pattern.

where we defined $\tilde{\phi}(\rho_\mathrm{T}^\mathrm{tot}) = \phi(\tilde{\rho}_\mathrm{M}(\rho_\mathrm{T}^\mathrm{tot}))$. As the production of the chemoattractant $\rho_\mathrm{C}$ increases with the local density of the thymocytes, this system allows for chemotaxis-driven accumulation of the thymocytes.[4]

To summarize the linear stability analysis, this mathematical analysis allows to determine the parameter region in which the model system spontaneously segregates into cortex and medulla. We then constructed two reduced mathematical models that either only show a reaction–diffusion or only a chemotaxis-driven instability. Using QSS approximations, we reduced these two model systems to two-component systems. The two-component reaction–diffusion system shows that the reaction–diffusion (Turing) instability is driven by the positive feedback in mTEC proliferation mediated by the activated thymocytes. In contrast, the two-component system for the chemotaxis-driven instability mechanism resembles the Keller–Segel model, clarifying that its instability is driven by the self-amplifying chemotactic aggregation of the thymocytes. In the full spatial thymic cross-talk model, both instability mechanisms play together, resulting in the increased unstable region of the full model (gray-shaded region) compared to the instability regions of the single mechanisms (blue- and orange-shaded region) shown in Fig. 4($A$) in the main text.

#### III. NUMERICAL SIMULATION AND ANALYSIS

In the main text, we make use of several observables to quantify the system's behavior in numerical simulations, such as the medullary volume fraction and the length scale of the cortex-medulla pattern. In the following, we first give details on the numerical simulations and then define the observables used.

##### A. Numerical simulation

The model simulations are performed using the time-dependent finite-element solver of Comsol Multiphysics (Version 6.0) [90] on a free tetrahedral (triangular) mesh with linear Lagrange elements of a maximal mesh-element size of 60µm in three, and 10µm in two dimensions. Linear stability analysis was performed using Wolfram Mathematica, and analysis was performed using Python 3.6, if not indicated otherwise. Model parameters are used as specified in Tables S1, S2.

[4] Also, note that the reaction terms give rise to a Jacobian with the signs

$$\begin{pmatrix} - & 0 \\ + & - \end{pmatrix}, \tag{38}$$

which does not fulfill the Segel–Jackson criterion and thus does not allow for a reaction–diffusion instability.

##### B. Observables

###### 1. Medullary volume fraction

One observable describing the medullary morphology is the medullary volume fraction $\varphi_\mathrm{M}(\boldsymbol{x})$ (see Figs. 3, 5$A$ in the main text). The medullary volume fraction is defined as the volume of the medulla divided by the volume of the full simulation domain. To determine its value, we define the (logistic) indicator function

$$\psi(\boldsymbol{x}) = \frac{1}{1 + e^{-(\rho_\mathrm{M}(\boldsymbol{x})-\mu)/10^{-6}\mathrm{µm}^{-3}}}\,. \tag{39}$$

The indicator function is close to zero for mTEC densities below the threshold $\rho_\mathrm{M}(\boldsymbol{x}) \ll \mu$ and rapidly increases to one around $\rho_\mathrm{M}(\boldsymbol{x}) = \mu = 10^{-4}\mathrm{µm}^{-3}$. The small density scale $10^{-6}\mathrm{µm}^{-3}$ in the exponent ensures a sharp transition between zero and one.

We average this indicator function over the simulation volume using Comsol's "Volume Average" (derived value) function. This function implements the spatial average

$$\langle \bullet \rangle \equiv \frac{1}{|\Omega|} \int_\Omega \mathrm{d}^d x \; \bullet \,. \tag{40}$$

Here, $\Omega$ is the two- ($d = 2$; used in Fig. 5$A$ in the main text) or three-dimensional ($d = 3$; used in Fig. 3 in the main text) simulation domain with the volume $|\Omega|$. We then define the medullary volume fraction as $\varphi_\mathrm{M} = \langle \psi \rangle$.

###### 2. Pattern length scale

Moreover, the medullary pattern is quantified by its typical pattern wavelength in Fig. 5$B$ in the main text. The pattern length scale is calculated using the connected two-point correlator for the mTEC density,

$$C(r) = \langle \rho_\mathrm{M}(0) \cdot \rho_\mathrm{M} \rangle - \langle \rho_\mathrm{M} \rangle^2\,, \tag{41}$$

using the average Eq. (40) calculated in Python (with a different normalization that will be irrelevant for the following).

From the radial correlation function $C(r)$, we determine its first minimum, whose value $r_0$ of the distance coordinate $r$ gives the typical length scale on which the pattern is anti-correlated. Thus, $r_0$ can be interpreted as half of the typical pattern length scale. To account for noise in the data that might produce local minima in an otherwise falling function, we determine the minimum by the following algorithm. We search for the minimum in a sliding window of 11 data points which we iteratively center around the newly found minimum. Once the minimum of the window is in its center, the iteration stops. We then determine the minimum by fitting a quadratic function to the 11 points in the window. If no minimum is found using this method, the length scale is set to the length associated with the correlator's absolute minimum.

###### 3. Thymocyte localization

Fig. 4*D, E* in the main text analyzes the localization of the thymocytes in the medullary structure. Specifically, we measure the local density of thymocytes within the medulla. Therefore, we define the average thymocyte density in the medulla $\langle\rho_\mathrm{T}(\boldsymbol{x})\rangle_\mathrm{M}$ as

$$\langle\rho_\mathrm{T}(\boldsymbol{x})\rangle_\mathrm{M} \equiv \frac{\langle\psi(\boldsymbol{x})\rho_T(\boldsymbol{x})\rangle}{\langle\psi(\boldsymbol{x})\rangle} \,. \tag{42}$$

Moreover, the average thymocyte density in the cortex $\langle\rho_\mathrm{T}(\boldsymbol{x})\rangle_\mathrm{C}$ is defined as

$$\langle\rho_\mathrm{T}(\boldsymbol{x})\rangle_\mathrm{C} \equiv \frac{\langle(1-\psi(\boldsymbol{x}))\rho_T(\boldsymbol{x})\rangle}{\langle(1-\psi(\boldsymbol{x}))\rangle} \,. \tag{43}$$

The values of these two averages in the simulations are plotted as bar charts in Fig. 4*D, E* in the main text.

##### C. Effective recognition rate

In Eq. (6) of the main text, we defined the effective recognition rate as

$$k_\mathrm{rec} = \frac{\langle k_\mathrm{r}\rho_\mathrm{T}\rho_\mathrm{M}\rangle}{\langle\rho_\mathrm{T}\rangle} \,. \tag{44}$$

We obtain this rate in simulations using the spatial average Eq. (40). We also note that for the uniform, non-segregated system, the effective recognition rate becomes

$$k_\mathrm{rec}^\mathrm{uniform} = k_\mathrm{r}\rho_\mathrm{M} \,. \tag{45}$$

#### IV. THYMOCYTE LOCALIZATION AND CHEMOTAXIS IN THE MEDULLARY PATTERN

In this section, we present additional results on self-reactive and self-tolerant thymocyte localization, as well as chemotaxis strength in dependence on the medulla-cortex pattern length scale.

##### A. Intramedullary thymocyte localization

As outlined in the main text, Sec. III B, chemotaxis drives self–reactive thymocytes to aggregate inside the medulla. In Fig. 4($D$), ($E$) of the main text, we show the average thymocyte densities in the cortex and medulla. However, the two thymocyte species show subtle differences in their distributions within the medulla, as shown in Fig. S3. The concentration of self–reactive thymocytes is highest at the edges of the medulla, and falls off toward the middle, as they are degraded inside the medulla. On the other hand, self–tolerant thymocytes are distributed roughly uniformly throughout the medulla because they are not negatively selected and degraded, and the rate of export into the body is much lower than the degradation rate of self-reactive thymocytes. Should this result hold in the experimental setting, we expect that it has downstream effects on the intra-medullary architecture, for example on distributions of medullary cells that rely on signals from activated thymocytes such as Aire$^+$ mTECs [33, 48].

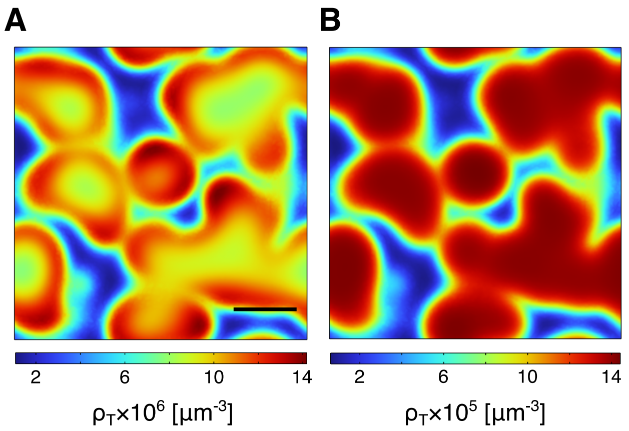


Figure S3. **Intra-medullary localization of self-reactive and tolerant thymocytes.** ($A$) Self-reactive thymocyte densities are highest at the medulla edge (cortico-medullary junction, CMJ). This is due to thymocyte flow into the medulla from the edge and degradation in the interior. ($B$) Self-tolerant thymocytes reach the highest density in the interior of the medulla, as their time scale of degradation due to thymus exit is far slower than the time it takes to migrate into the medulla (see also Sec. I B 16). C.f. also Fig. 4($C$), main text. Scale bar 500µm.

##### B. Dependence of chemotaxis on the medullary pattern length scale

As discussed in Sec. III D of the main text, the length scale of the cortex-medulla pattern impacts the effective recognition rate of self–reactive thymocytes. We relate the increase in the effective recognition rate to a faster transition of the thymocytes from the cortex into the medulla. Here, we measure the average chemotactic speed $\langle|\boldsymbol{v}_\mathrm{C}|\rangle = \langle T\chi(\rho_\mathrm{C})\,|\nabla\rho_\mathrm{C}|\rangle$ of the thymocytes in the simulations. Figure S4 shows that the chemotaxis speed is strongly dependent on the length scale of the cortex-medulla pattern. It is small for small pattern length scales (around 200µm), then increases toward intermediate length scales, before falling off at length scale $\gtrsim$ 1mm. The fall-off in the chemotaxis speed at large pattern length scales coincides with the fall-off in the effective recognition rate [cf. Fig. 6($C$) in the main text]. The chemotaxis speed also falls off toward short pattern length scales while the effective recognition rate levels off towards the value of homogeneously distributed mTECs for short length–scales (Fig. 6($C$), main text). The reason for this difference at short length scales is that chemotaxis becomes less important for the recognition process as the thymocyte densities are smeared out by diffusion.

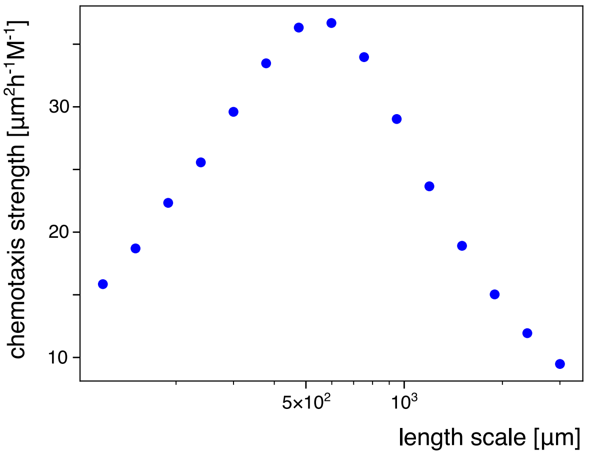


Figure S4. **Medullary length scale modulates chemotaxis speed.** The averaged chemotaxis speed $\langle|\boldsymbol{v}_{\mathrm{C}}|\rangle = \langle T\chi(\rho_{\mathrm{C}})\,|\nabla\rho_{\mathrm{C}}|\rangle$ is largest for intermediate length scales of medullary lamellae. The chemotaxis speed peaks at the same medullary length scale [c.f. Fig. 6($C$), main text] as selection efficiency.

[10] J.-E. Park, R. A. Botting, C. Domínguez Conde, D.-M. Popescu, M. Lavaert, D. J. Kunz, I. Goh, E. Stephenson, R. Ragazzini, E. Tuck, A. Wilbrey-Clark, K. Roberts, V. R. Kedlian, J. R. Ferdinand, X. He, S. Webb, D. Maunder, N. Vandamme, K. T. Mahbubani, K. Polanski, L. Mamanova, L. Bolt, D. Crossland, F. de Rita, A. Fuller, A. Filby, G. Reynolds, D. Dixon, K. Saeb-Parsy, S. Lisgo, D. Henderson, R. Vento-Tormo, O. A. Bayraktar, R. A. Barker, K. B. Meyer, Y. Saeys, P. Bonfanti, S. Behjati, M. R. Clatworthy, T. Taghon, M. Haniffa, and S. A. Teichmann, A cell atlas of human thymic development defines T cell repertoire formation, Science **367**, eaay3224 (2020).

[26] P. Brennecke, A. Reyes, S. Pinto, K. Rattay, M. Nguyen, R. Küchler, W. Huber, B. Kyewski, and L. M. Steinmetz, Single-cell transcriptome analysis reveals coordinated ectopic gene-expression patterns in medullary thymic ep-

ithelial cells, Nature Immunology **16**, 933 (2015).
[27] C. Bornstein, S. Nevo, A. Giladi, N. Kadouri, M. Pouzolles, F. Gerbe, E. David, A. Machado, A. Chuprin, B. Tóth, O. Goldberg, S. Itzkovitz, N. Taylor, P. Jay, V. S. Zimmermann, J. Abramson, and I. Amit, Single-cell mapping of the thymic stroma identifies IL-25-producing tuft epithelial cells, Nature **559**, 622 (2018).
[28] J. Baran-Gale, M. D. Morgan, S. Maio, F. Dhalla, I. Calvo-Asensio, M. E. Deadman, A. E. Handel, A. Maynard, S. Chen, F. Green, R. V. Sit, N. F. Neff, S. Darmanis, W. Tan, A. P. May, J. C. Marioni, C. P. Ponting, and G. A. Holländer, Ageing compromises mouse thymus function and remodels epithelial cell differentiation, eLife **9**, e56221 (2020).
[29] K. L. Wells, C. N. Miller, A. R. Gschwind, W. Wei, J. D. Phipps, M. S. Anderson, and L. M. Steinmetz, Combined transient ablation and single-cell RNA-sequencing reveals the development of medullary thymic epithelial cells, eLife **9**, e60188 (2020).
[30] E. W. Shores, W. Van Ewijk, and A. Singer, Disorganization and restoration of thymic medullary epithelial cells in T cell receptor-negative scid mice: Evidence that receptor-bearing lymphocytes influence maturation of the thymic microenvironment, European Journal of Immunology **21**, 1657 (1991).
[31] M. Irla, S. Hugues, J. Gill, T. Nitta, Y. Hikosaka, I. R. Williams, F.-X. Hubert, H. S. Scott, Y. Takahama, G. A. Holländer, and W. Reith, Autoantigen-Specific Interactions with CD4+ Thymocytes Control Mature Medullary Thymic Epithelial Cell Cellularity, Immunity **29**, 451 (2008).
[32] M. Irla, L. Guerri, J. Guenot, A. Sergé, O. Lantz, A. Liston, B. A. Imhof, E. Palmer, and W. Reith, Antigen Recognition By Autoreactive Cd4+ Thymocytes Drives Homeostasis Of The Thymic Medulla, PLoS ONE **7**, e52591 (2012).
[33] N. Lopes, N. Boucherit, J. C. Santamaria, N. Provin, J. Charaix, P. Ferrier, M. Giraud, and M. Irla, Thymocytes trigger self-antigen-controlling pathways in immature medullary thymic epithelial stages, eLife **11**, e69982 (2022).
[34] L. Klein, B. Kyewski, P. M. Allen, and K. A. Hogquist, Positive and negative selection of the T cell repertoire: What thymocytes see (and don't see), Nature Reviews Immunology **14**, 377 (2014).
[35] M. S. Anderson, E. S. Venanzi, L. Klein, Z. Chen, S. P. Berzins, S. J. Turley, H. von Boehmer, R. Bronson, A. Dierich, C. Benoist, and D. Mathis, Projection of an Immunological Self Shadow Within the Thymus by the Aire Protein, Science **298**, 1395 (2002).
[36] L. Onder, V. Nindl, E. Scandella, Q. Chai, H.-W. Cheng, S. Caviezel-Firner, M. Novkovic, D. Bomze, R. Maier, F. Mair, B. Ledermann, B. Becher, A. Waisman, and B. Ludewig, Alternative NF-$\kappa$B signaling regulates mTEC differentiation from podoplanin-expressing precursors in the cortico-medullary junction, European Journal of Immunology **45**, 2218 (2015).
[37] G. Anderson and Y. Takahama, Thymic epithelial cells: Working class heroes for T cell development and repertoire selection, Trends in Immunology **33**, 256 (2012).
[38] H. Takaba, Y. Morishita, Y. Tomofuji, L. Danks, T. Nitta, N. Komatsu, T. Kodama, and H. Takayanagi, Fezf2 Orchestrates a Thymic Program of Self-Antigen Expression for Immune Tolerance, Cell **163**, 975 (2015).
[39] K. M. Ashby and K. A. Hogquist, A guide to thymic selection of T cells, Nature Reviews Immunology , 1 (2023).
[40] E. R. Breed, M. Watanabe, and K. A. Hogquist, Measuring Thymic Clonal Deletion at the Population Level, The Journal of Immunology **202**, 3226 (2019).
[41] J. N. Lancaster, H. M. Thyagarajan, J. Srinivasan, Y. Li, Z. Hu, and L. I. R. Ehrlich, Live-cell imaging reveals the relative contributions of antigen-presenting cell subsets to thymic central tolerance, Nature Communications **10**, 2220 (2019).
[42] J. P. van Meerwijk, S. Marguerat, R. K. Lees, R. N. Germain, B. Fowlkes, and H. R. MacDonald, Quantitative Impact of Thymic Clonal Deletion on the T Cell Repertoire, Journal of Experimental Medicine **185**, 377 (1997).
[43] M. Hinterberger, M. Aichinger, O. Prazeres da Costa, D. Voehringer, R. Hoffmann, and L. Klein, Autonomous role of medullary thymic epithelial cells in central CD4+ T cell tolerance, Nature Immunology **11**, 512 (2010).
[44] J. S. Perry, C.-W. J. Lio, A. L. Kau, K. Nutsch, Z. Yang, J. I. Gordon, K. M. Murphy, and C.-S. Hsieh, Distinct Contributions of Aire and Antigen-Presenting-Cell Subsets to the Generation of Self-Tolerance in the Thymus, Immunity **41**, 414 (2014).
[45] H. Wang and J. C. Zúñiga-Pflücker, Thymic Microenvironment: Interactions Between Innate Immune Cells and Developing Thymocytes, Frontiers in Immunology **13**, 10.3389/fimmu.2022.885280 (2022).
[46] N. Lopes, A. Sergé, P. Ferrier, and M. Irla, Thymic Crosstalk Coordinates Medulla Organization and T-Cell Tolerance Induction, Frontiers in Immunology **6**, 10.3389/fimmu.2015.00365 (2015).
[47] D. Malhotra, J. L. Linehan, T. Dileepan, Y. J. Lee, W. E. Purtha, J. V. Lu, R. W. Nelson, B. T. Fife, H. T. Orr, M. S. Anderson, K. A. Hogquist, and M. K. Jenkins, Tolerance is established in polyclonal CD4+ T cells by distinct mechanisms, according to self-peptide expression patterns, Nature Immunology **17**, 187 (2016).
[48] M. Irla, J. Guenot, G. Sealy, W. Reith, B. A. Imhof, and A. Sergé, Three-Dimensional Visualization of the Mouse Thymus Organization in Health and Immunodeficiency, The Journal of Immunology **190**, 586 (2013).
[49] M. Sakata, I. Ohigashi, and Y. Takahama, Cellularity of Thymic Epithelial Cells in the Postnatal Mouse, The Journal of Immunology **200**, 1382 (2018).
[50] S. R. Daley, D. Y. Hu, and C. C. Goodnow, Helios marks strongly autoreactive CD4+ T cells in two major waves of thymic deletion distinguished by induction of PD-1 or NF-$\kappa$B, Journal of Experimental Medicine **210**, 269 (2013).
[51] G. L. Stritesky, Y. Xing, J. R. Erickson, L. A. Kalekar, X. Wang, D. L. Mueller, S. C. Jameson, and K. A. Hogquist, Murine thymic selection quantified using a unique method to capture deleted T cells, Proceedings of the National Academy of Sciences **110**, 4679 (2013).
[52] I. L. Dzhagalov, K. G. Chen, P. Herzmark, and E. A. Robey, Elimination of Self-Reactive T Cells in the Thymus: A Timeline for Negative Selection, PLoS Biology **11**, e1001566 (2013).
[53] N. S. Kurd, L. K. Lutes, J. Yoon, S. W. Chan, I. L. Dzhagalov, A. R. Hoover, and E. A. Robey, A role for phagocytosis in inducing cell death during thymocyte negative selection, eLife **8**, e48097 (2019).

[90] COMSOL Multiphysics, COMSOL AB (2019).
